# Cryptic endogenous retrovirus subfamilies in the primate lineage

**DOI:** 10.1101/2023.12.07.570592

**Authors:** Xun Chen, Zicong Zhang, Yizhi Yan, Clement Goubert, Guillaume Bourque, Fumitaka Inoue

## Abstract

Many endogenous retroviruses (ERVs) in the human genome are primate-specific and have contributed novel cis-regulatory elements and transcripts. However, current approaches for classifying and annotating ERVs and their long terminal repeats (LTRs) have limited resolution and are inaccurate. Here, we developed a new annotation based on phylogenetic analysis and cross-species conservation. Focusing on the evolutionary young MER11A/B/C subfamilies, we revealed the presence of 4 ‘new subfamilies’, that better explained the epigenetic heterogeneity observed within the MER11 instances, suggesting a new annotation for 412 (19.8%) of these repeat elements. Furthermore, we functionally validated the regulatory potential of these four new subfamilies using a massively parallel reporter assay (MPRA), which also identified motifs associated with their differential activities. Combining MPRA with new annotations across primates revealed an apes-specific gain of SOX related motifs through a single-nucleotide deletion. Lastly, by applying our approach across 53 simian-enriched LTR subfamilies, we defined a total of 75 new subfamilies and found that 3,807 (30.0%) instances from 26 LTR subfamilies could be categorized into a novel annotation, many of which with a distinct epigenetic profile. Thus, with our refined annotation of simian-enriched LTRs, it will be possible to better understand the evolution in primate genomes and potentially identify new roles for ERVs and their LTRs in the hosts.

## INTRODUCTION

Transposable elements (TEs) occupy nearly half of the human genome, and recent genomic and epigenomic analyses have revealed that many have been co-opted by the host(*1–6*). In particular, at least 8% of the human genome finds its origin in endogenous retroviruses (ERVs)(*7*, *8*), which includes subfamilies of ERV internal and long terminal repeat (LTR) sequences separately, while LTR sequences are often found in open chromatin and have the potential to act as genomic regulatory elements(*1*, *2*, *4*, *9*). ERVs originate from retrovirus infections, where the viral fragment interacts with transcription factors (TFs) in the host cell via its long terminal repeat (LTR) sequences to express viral RNA, then spreads throughout the genome by a copy-and-paste mechanism(*10*). ERVs including their internal and LTR sequences could spread in the host genome mainly through the vertical and horizontal transmission during evolution(*11–13*). To limit the deleterious effects of uncontrolled transposition, host cells have evolved multiple defense mechanisms to silence ERVs including CpG methylation, m6A RNA methylation, RNA interference (PIWI-associated small RNA), KRAB-associated repressors genes and H3K9me3 modification(*14–16*). Along with this active silencing, most TEs, including LTR sequences, accumulate mutations, which eventually result in their inactivation(*17*). Nevertheless, some LTR sequences may retain their regulatory activity or acquire beneficial mutations within transcription factor binding motifs(*6*). This can impact the regulatory activity of nearby gene expression, and contribute to the adaptation of the LTR copies in the host genome(*6*, *18*, *19*). Additionally, ERVs have a higher chance of survival in the next generation when they are expressed in germline or pluripotent cells during early embryonic development(*20–22*). In fact, several LTR subfamilies contain binding sequences for pluripotency transcription factors such as POU5F1 and SOX2(*6*, *23*).

Many ERVs integrated into the primate genomes after the divergence from other mammals(*24*). In the process, these relatively young ERV elements have contributed a substantial number of regulatory sequences to the human genome(*9*) and are associated with the evolution of TF binding sites during primate evolution(*25*). Some subfamilies retain regulatory or transcriptional activity and have influenced human transcriptional networks(*1*, *2*, *4*, *26–30*). HERVH-LTR7, for instance, is considered endogenized in the primate lineage, since many of its instances (copies) have been co-opted by the host as gene regulatory elements in pluripotent stem cells(*30*). LTR5_Hs has also been shown, using a chimeric array of gRNA oligos (CARGO) and CRISPR, to act as enhancers that regulate hundreds of human genes(*29*). Furthermore, the divergent expansion of LTR subfamilies within individual branches of the primate lineage, which provides a wealth of species-specific enhancers, significantly influences the regulatory network of these genomes during speciation(*31*).

The proper classification and annotation of LTR instances is critical to understand their evolution and potential impact on the host(*32*). The standard approach used to characterize TE instances relies on homology between genomic sequences and curated TE libraries, and aims to attribute a unique family or subfamily name to a group of monophyletic sequences(*33*). Although an important effort of manual curation has been applied to the TEs in the human lineage, a correct classification and annotation of these repeat elements remains a challenging problem(*34–37*). Deep analysis of ERVs/LTRs across primates further emphasize the complexity of their evolution and potential annotation problem(*38–40*). Historical nomenclature discrepancy, a high degree of sequence similarity between related yet distinct monophyletic groups, as well as recombination events within ERVs or between ERVs and exogenous viruses has led to various misclassifications(*41*). Furthermore, instances of an established ERV internal and LTR subfamily continuously evolve further into divergent sub-lineages, introducing another layer of complexity(*42*, *43*). Phylogenetic analyses have been used to study ERV/LTR sequence evolution with either a small set of full length ERV or solo LTR sequences(*44*, *45*), or to interpret their regulatory heterogeneity(*46*). Unfortunately, most of these studies relied on the current ERV internal and LTR classification. More recently, phylo-regulatory approaches, combining phylogenetic reconstruction of LTR subfamilies and layering of epigenetic data, have helped overcoming some of these issues for specific LTR subfamilies, e.g., LTR7(*42*).

Considering the nature of LTRs, we hypothesized that the proper classification and annotation of LTR copies would require investigating their phylogenetic relationships in order to infer their evolution and function. In this work, we present an improved annotation of simian-enriched LTR subfamilies in the human genome using a phylogenetic approach that combines sequence analysis and the presence of orthologous instances in other species.

## RESULTS

### The LTR subfamilies spreading in the primate lineage are heterogenous

We wanted to investigate the evolution of LTR subfamilies (i.e., subfamilies of LTR sequences but not ERV internal sequences) and selected 179 subfamilies enriched in the human genome with copies in the marmoset or more closely related primate species, but mostly absent from lemur using the liftOver approach (hence putatively integrated or extensively expanded since the simian ancestor ∼40 million years ago) (Fig. 1a). The selected subfamilies were consistently observed to be enriched in simian genomes by further examining a total of 47 primate species (see Methods, Supplementary Fig. 1a and Supplementary Data 1). We focused on LTR subfamilies since they usually contain most of the sequences driving ERVs regulatory and transcriptional activity. Among them, we further selected 35 subfamilies with limited shared repeat instances (≤ 60%) in the macaque genome (Supplementary Fig. 1b). As expected, the use of newer genome builds may improve their locations in larger contigs but not significantly change the presence/absence calls (Supplementary Fig. 1c and Supplementary Data 2). Based on a network analysis of repeat consensus sequence similarity(*47*), we also identified 26 closely related LTR subfamilies for a total of 61 putative simian-enriched subfamilies organized in 19 groups, which refers to the subfamilies that were either first integrated or widely spread in simian genomes (Supplementary Fig. 2 and Supplementary Fig. 3). For instance, MER9a1, MER9a2, MER9a3, and MER9B were clustered together and were distinct from other subfamilies. Next, by examining the distribution of divergence rates of instances within the 61 LTR subfamilies (Fig. 1b and Supplementary Fig. 4a), we found that 28 (46%) had a non-normal distribution (chi-square *p* value ≤ 0.001), including LTR12B, LTR5B, LTR7Y, LTR61, LTR5_Hs and others showing a bimodal distribution. This is consistent with a recent report characterizing multiple subgroups within the LTR7 subfamilies(*42*), and suggests that many simian-enriched LTR subfamilies are heterogeneous.

**Figure 1.**
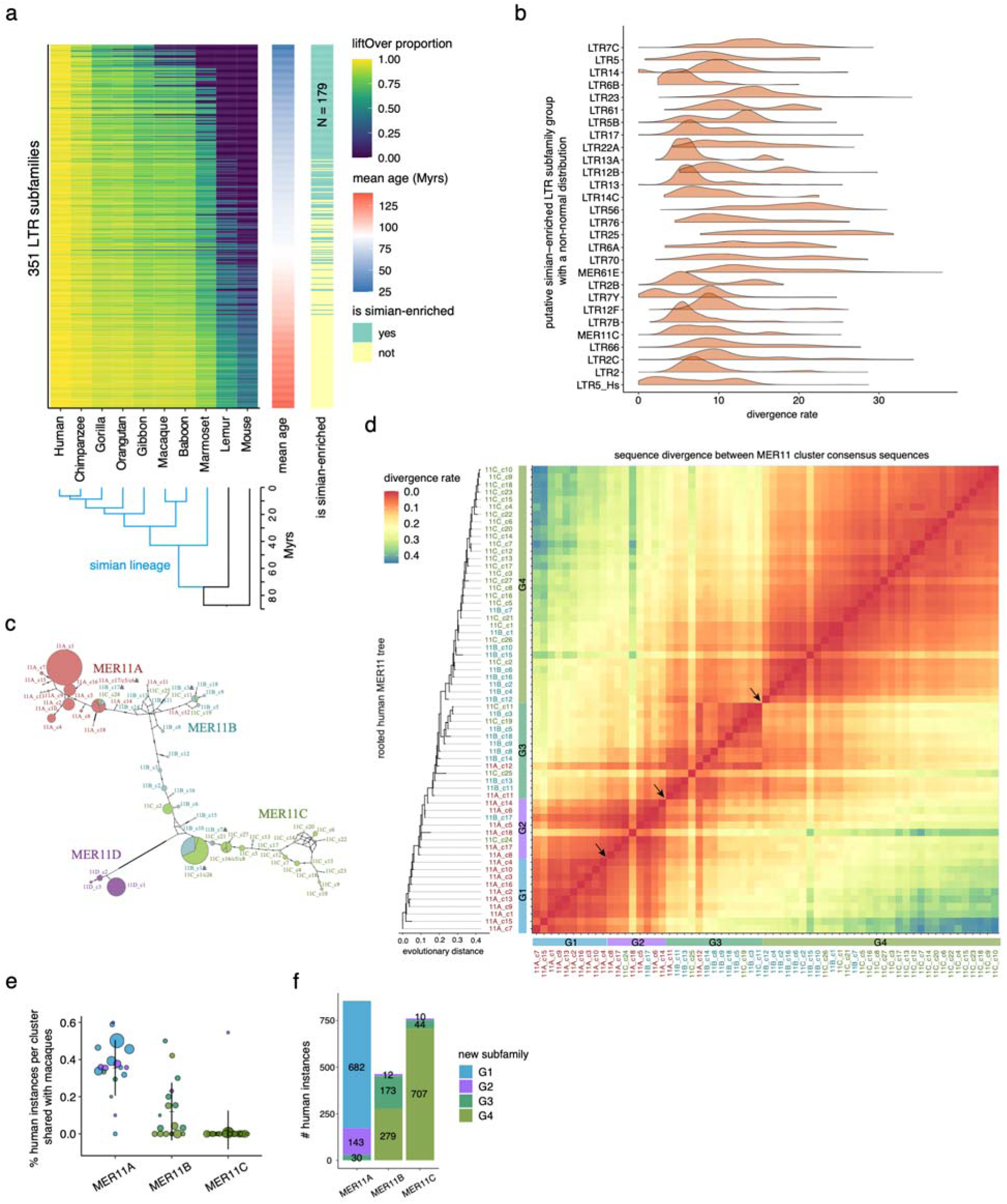
The sequence heterogeneity of simian-enriched LTR subfamilies. **a** LiftOver analysis of LTR subfamilies (not including ERV internal sequences) from human (hg19) to other primate and mouse genomes (see Methods). Evolutionary ages were estimated based on the divergence rates. LTR subfamilies that have a minimum of 100 instances (≥ 200 bp) and a maximum of 20% of instances shared with the lemur genome were selected. **b** Divergence rate (% substitutions) distribution of instances relative to the subfamily consensus sequence. Subfamilies that have unexpected distributions (Chi-square test, Bonferroni adjusted *p* values < 0.001) are shown. **c** Median-joining network of the 66 MER11 clusters. Circle sizes refer to the relative number of instances between clusters. Ticks indicate the number of nucleotide mutations between cluster consensus sequences. Clusters derived from different MER11 original subfamilies are in different colors. Pie charts indicate clusters that have the same consensus sequences after the removal of gaps. **d** Rooted tree containing 63 MER11A/B/C clusters. Heatmap displays divergence rates between each pair of cluster consensus sequences. Four new subfamilies are identified using different colors (see Methods). Arrows indicate the high divergence rates between clusters from adjacent new subfamilies. Clusters derived from each original subfamily are in different colors. **e** Proportion of human instances shared with macaques. Dot size refers to the number of instances per cluster. **f** Number of instances per MER11A/B/C original subfamily assigned to each new subfamily.

From this list, the MER11 subfamilies are of particular interest, because they were amongst the youngest, had the lowest proportion of shared instances in macaque and displayed non-normal divergence rate distributions (Fig. 1b, Supplementary Fig. 1b and Supplementary Fig. 4a-b). To explore the variability in this group, we built an unrooted tree separately for MER11A, MER11B and MER11C based on the multiple sequence alignment of all the repeat instances. Another MER11 subfamily, MER11D, was also analyzed to confirm its distal relationship. Following a method described previously(*42*), we grouped instances into 66 clusters based on internal branch length: 18 for MER11A (labeled 11A_c1-18), 18 for MER11B (labeled 11B_c1-18), 27 for MER11C (labeled 11C_c1-27) and 3 clusters for MER11D (labeled 11D_c1-3) (Supplementary Fig. 4c). To understand the relationship among the 66 MER11 clusters, we performed a median-joining network analysis using the cluster consensus sequences - a method used to infer intraspecific phylogenies(*48*) (Fig. 1c). As expected, MER11D clusters were grouped as an independent branch, and we found that most MER11A and MER11C clusters were grouped together. Notably, many MER11B clusters were dispersed between MER11A and MER11C clusters except 11B_c3/c5/c9/c18 that were found to be on a separate branch.

To further understand the evolution among these MER11A/B/C subfamilies, we inferred rooted trees using the non-reversible model without the use of an outgroup(*49*) (see Methods). After lifting over to other primate genomes the instances from each cluster, we selected the root that was most consistent with the proportions of shared instances across species (Supplementary Fig. 4d). This rooted tree was also found to be consistent with the network we constructed, further confirming the expansion progress of these repeat instances (Supplementary Fig. 4e). Based on the rooted tree and internal branch lengths between cluster consensus sequences, we defined four new subfamilies (see Methods, Fig. 1d), i.e., MER11_G1, MER11_G2, MER11_G3 and MER11_G4. As expected, instances in these new subfamilies displayed more homogenous divergence rates compared to the original subfamilies (Supplementary Fig. 3b and Supplementary Fig. 3f). Notably, some clusters of instances from the evolutionary old MER11A were put in MER11_G1; while others were grouped with clusters from MER11B/C to form MER11_G2. Moreover, half of MER11B clusters and two MER11A and three MER11C clusters were grouped into MER11_G3, and most MER11C clusters and another half of MER11B clusters were grouped into MER11_G4. We also found that clusters with a relative higher or lower liftOver rate to macaques compared to other clusters (e.g., 11B_c17, 11C_c27, and 11B_c1) were more likely to be reassigned to new subfamilies (Fig. 1e). Notably, if we were to select the top new subfamiliy to represent each original annotation, we found that a total of 412 (19.8%) MER11 instances would be annotated differently (Fig. 1f and Supplementary Data 3). Based on the new annotation, we further profile their distribution on the human genome. As expected, instances from different new subfamilies are overally randomly distributed across each genome with some enrichment in specific chromosomes, e.g., MER11_G1 in chr4 and MER11_G3 in chrY (Supplementary Fig. 4f). Taken together, detailed sequence analysis revealed four new subfamilies with many MER11 instances that contrasted with the original subfamily classification.

### MER11 new subfamilies display more consistent epigenetic profiles as compared to original MER11 subfamilies

Given that the new subfamilies of MER11 were more homogeneous based on divergence rates (Supplementary Fig. 4g), we hypothesized that they would also have more consistent epigenetic profiles in human cells. As endogenous retrovirus expression and endogenization often occurs in early developmental stages(*18*, *50*), we first compared chromatin accessibility and H3K27ac active histone mark in human embryonic stem cells (hESCs) and in hESC-derived neural progenitor cells (NPCs) using two published datasets(*51*, *52*). From this, we identified 18 LTR subfamilies that were significantly enriched in hESCs (Supplementary Fig. 5a-c). In particular, we found that MER11B and MER11C subfamilies showed relatively high cell-type specificity in hESCs and mesendoderm cells (Supplementary Fig. 5d). Next, we performed an integrative analysis combining the phylogenetic trees built from MER11A/B/C clusters and their epigenetic profiles in hESCs using a panel of 27 histone marks and 65 transcription factor (TF) ChIP-seq datasets from the ENCODE project (Fig. 2a). Notably, we observed that specific clusters within the subfamily trees were enriched for specific histone marks and TF peaks. For example, the younger MER11A clusters, with a long evolutionary distance to the root, were enriched for active histone marks and TF peaks. In contrast, the more ancient MER11B and MER11C clusters were enriched for active histone marks and TF peaks. The youngest MER11C clusters were enriched for active histone marks but less enriched for TF peaks. Thus, each original subfamily had a high epigenetic heterogeneity.

**Figure 2.**
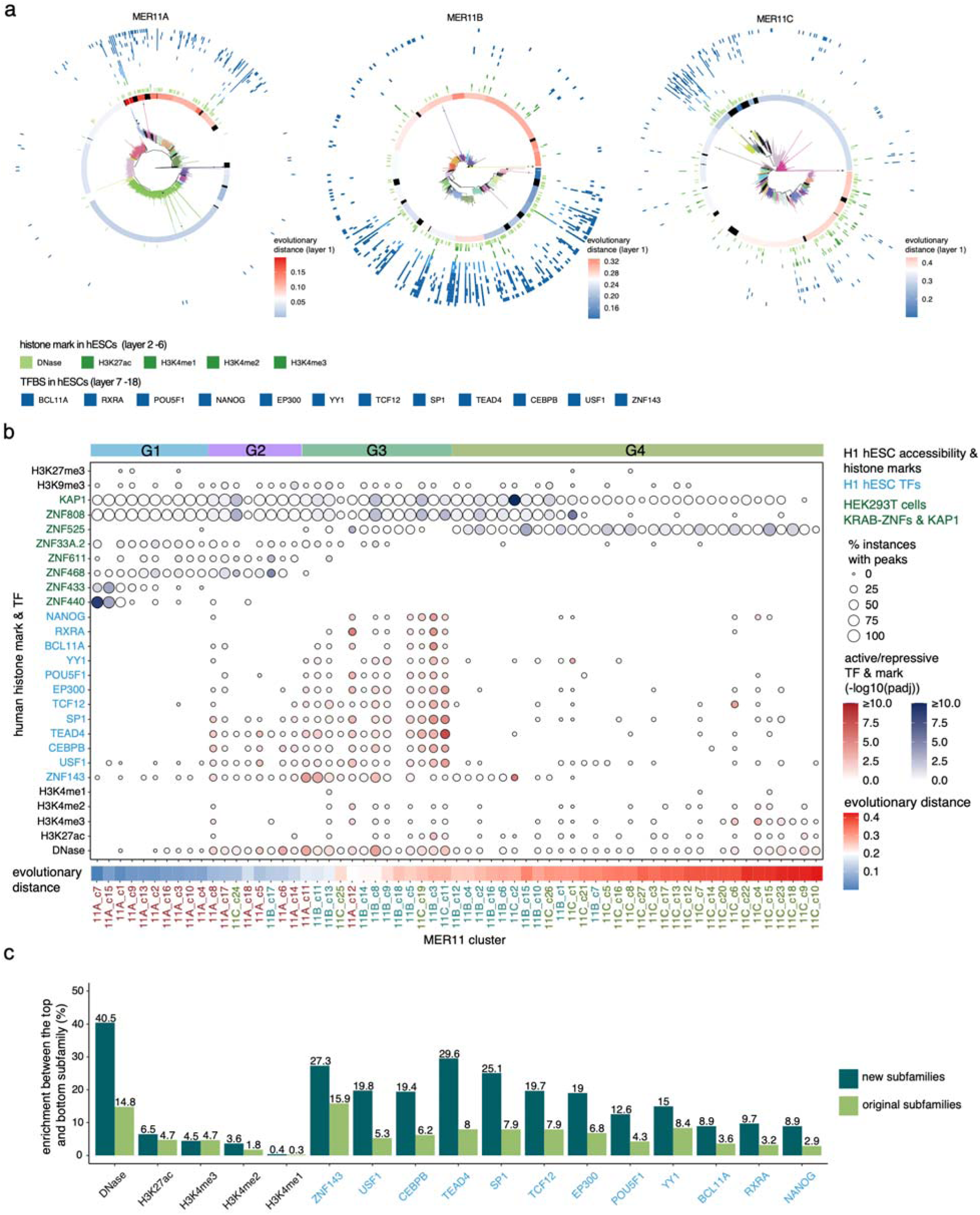
MER11 new subfamilies display more consistent epigenetic profiles as compared to original annotations. **a** Circular plots of MER11A/B and C. Unrooted trees are used to order instances (center). Evolutionary distance per cluster was calculated using the root from the tree of Fig. 1D. Active histone marks and significantly enriched TFBSs in any MER11 original subfamilies are shown (see Methods). **b** Enrichment of active histone marks and TF peaks for every MER11 cluster ordered based on the evolutionary distance from the root. Active and repressive histone marks and significantly enriched TFs and KRAB-ZNFs in any MER11 original subfamily are shown. New subfamilies are highlighted on the top and separated by dotted lines. Clusters derived from each original subfamily are colored differently. **c** Enrichment of active histone marks and TF peaks in new versus original subfamilies. Enrichment was computed as the proportion of peaks-associated instances in the top new subfamily, the one with the highest proportion of peak-associated instances, minus the proportion in the bottom new subfamily, the one with the lowest proportion, for each histone mark and TF (in blue). The same was calculated between the top original subfamily and the bottom original subfamily.

To inspect whether our four new subfamilies of MER11 displayed a more consistent epigenetic profile compared to the original subfamily annotations, we rearranged the epigenetic data, using the rooted tree defined in the previous section, and observed distinct patterns of epigenetic states between new subfamilies (Fig. 2b). For instance, MER11_G1 lacked TF peaks and active histone marks; MER11_G3 was significantly enriched for open chromatin and most TF peaks, followed by MER11_G2; Among MER11_G4 clusters, only the youngest ones were enriched for active histone marks. Because of the balance between chromatin accessibility and repression over TEs(*53*), we also looked at the enrichment of KRAB-ZNFs and KAP1 (Supplementary Fig. 5e-f) binding in HEK293T cells. We further observed the sequential loss of ZNF440, ZNF433, ZNF468, ZNF611, ZNF33A, ZNF808 binding and the gain of ZNF525 binding along the evolution of the MER11 clusters (Fig. 2**b**). KAP1 binding was mostly enriched in MER11_G3 and relatively old MER11_G4 clusters, which was consistent with the enrichment of ZNF808 binding. Finally, compared to the original annotation of the MER11 subfamilies, we found that the four new subfamilies achieved a higher specificity across these active marks (Fig. 2c and Supplementary Fig. 5g). For example, TEAD4 enrichment was 29.6% using new subfamilies (it overlapped 29.6% of instances in MER11_G3 but 0% of instances in MER11_G1) which is much higher than original subfamily annotations (the enrichment was 8% since it overlapped 11% of MER11B instances but 3% instances in MER11A). Taken together, the four new subfamilies of MER11 appear to resolve the epigenetic heterogeneity within MER11 instances, and we found that relative age was associated with distinct regulatory profiles.

### MPRA confirms the regulatory potential of MER11 new subfamilies and reveals associated TF binding motifs

To further assess the biological relevance of the reconstituted MER11 subfamily annotations, we leveraged a massively parallel reporter assay (MPRA). First, we identified two peaks of accessible regions within the MER11A/B/C instances, and extracted their sequences (∼250-bp) as the frames - putative functional sequences of a suitable length - to be analyzed by MPRA (Fig. 3a and Supplementary Fig. 6a-b). As controls, we analyzed two older LTR subfamilies, MER34 and MER52 (Fig. 3b); MER34A1/C_ subfamilies were enriched for both ATAC-seq and H3K27ac peaks in hESCs compared to NPCs, while the enrichment of MER52C subfamily was slightly increased during the NPC differentiation (Supplementary Fig. 5b-c). We then retrieved homologous sequences in the human, chimpanzee and macaque genomes to characterize the regulatory potential of a large fraction of the observed evolutionary variants (Supplementary Fig. 6c). Some sequences had to be excluded due to the high number of mutations and truncations relative to the core frames used for the MPRA experiment (Methods). In the end, we analyzed 16,929 unique LTR sequences, including 6,912 MER11s, 5,751 MER34s, and 4,266 MER52s sequences, together with 100 positive and 100 negative control sequences (Fig. 3c and Supplementary Data 3-4), in both human iPSCs and iPSC-derived NPCs and in triplicates. We analyzed only high-quality sequences that were observed with at least 10 barcodes associated in the library (> 80% of MER11/52 and 30-75% of MER34, Supplementary Fig. 6d). Lower quality observed for MER34 was probably due to low GC content and long insert length (data not shown). The RNA/DNA ratios between replicates were strongly correlated (*R^2^* = 0.83), including for the positive and negative controls, indicating the accuracy of the MPRA measurements (Supplementary Fig. 6e).

**Figure 3.**
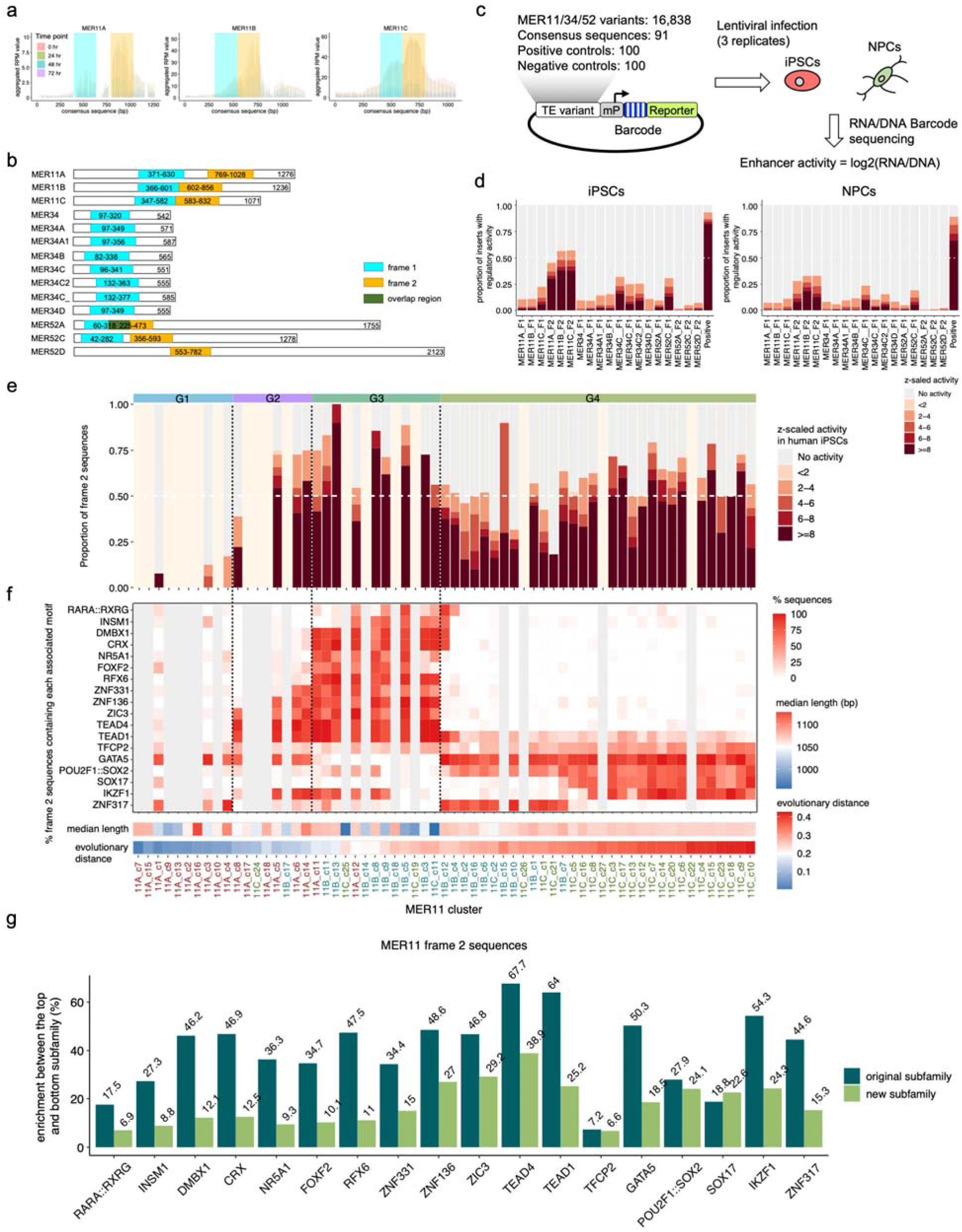
MPRA and MER11 new annotations help resolve the functional heterogeneity of MER11 subfamilies. **a** Determination of accessible regions along the consensus sequence per subfamily. ATAC-seq RPM (reads per million) values are aggregated across instances. **b** MER11, MER34, and MER52 consensus sequence frames designed for measuring enhancer activity using MPRA. **c** MPRA experiment workflow. MER11/34/52 variants, consensus, and control sequences were inserted into the MPRA vector with random barcodes. The MPRA library was infected into iPSCs or NPCs using lentivirus with three replicates. RNA and DNA barcodes were measured to quantify their enhancer activity. **d** Proportion of active sequences per sequence frame. Chimpanzee and macaque sequences orthologous to each cluster are also included. The normalization of MPRA activity was described in Methods. **e** Proportion of active MER11 frame 2 sequences per cluster. Clusters with less than 10 instances measured by MPRA are not shown. Background is in light pink color. Chimpanzee and macaque frame 2 sequences orthologous to each cluster are also included. **f** Percentage of associated motifs in the frame 2 sequences across MER11 clusters and new subfamilies. Chimpanzee and macaque sequences are also included. Clusters derived from each original subfamily are colored differently. **g** Motif enrichment in new versus original subfamilies. Enrichment was computed as the proportion of instances containing each motif in the top new subfamily, the one with the highest proportion, minus the bottom new subfamily, the one with the lowest proportion. Motif enrichment was computed similarly between the top and the bottom original subfamily.

After normalization, we found that MER11 frame 2 sequences showed higher MPRA activities compared to frame 1 sequences, which was consistent with the chromatin accessibility data (Fig. 3d and Supplementary Fig. 6f and Supplementary Data 5). Specifically, half of the MER11 frame 2 sequences were highly active (z-scaled activity ≥ 2) in human iPSCs, while the proportions stayed around 10% across MER11 frame 1 sequences. We next examined the MPRA activity among the human MER11 clusters and new subfamilies. Notably, even though the overall activity of frame 1 remained low, the activity levels were increased for the more recent MER11_G4 clusters (Supplementary Fig. 6g). For frame 2, the activity varied between new subfamilies, specifically some clusters from MER11_G2/3 and young clusters from MER11_G4 had the highest activity levels (Fig. 3e, Supplementary Data 3 and Supplementary Data 5). Most instances from the oldest MER11_G1 new subfamily, could not be analyzed by MPRA due to mutations and truncations; however, among the remaining sequences, the reported activity levels were low.

One of the reasons for generating MPRA data was to implement a TE-wide motif association analysis approach to identify TF binding motifs contributing to the activity detected(*54*). Among all frame 1 genomic sequences, we identified SP3 and related motifs (e.g., KLF12) followed by ZICs, POU::SOXs and SOXs motifs (Supplementary Fig. 7). We then looked at the MER11 frame 2 sequences and identified many motifs that were significantly associated with the MPRA activity including SOXs, POU2F1::SOX2, PITXs, ZICs, and TEADs. The motifs found to be enriched in MER34 (e.g., SPs and POU::SOXs) and MER52 (e.g., SPs, KLFs) were quite different. As expected, many TF motifs were strongly correlated with each other, such as RFX6, DMBX1, FOXF2, NR5A1, INSM1, and RARA::RXRG in MER11 frame 2 sequences (Supplementary Fig. 8). For each motif group, we only kept the most strongly associated motif (*p* ≤ 1 × 10^−10^). Next, we looked at the proportion of MER11 frame 2 sequences containing each motif (Fig. 3f). Notably, we observed a clear clustering of most motifs amongst the different new subfamilies. For instance, we observed the unique enrichment of NR5A1, FOXF2, and RFX6 related motifs in MER11_G3; ZNF136, ZIC3, and TEAD4 in both MER11_G3/G4; POU2F1::SOX2 and SOX17 in MER11_G4.

Finally, we investigated whether these TF motifs overlapped with nucleotides associated with the MPRA activity based on TE-wide nucleotide association analysis(*55*). Focusing on MER11 frame 2, we identified 15 single nucleotide variants and 32 indels that were significantly associated with the MPRA activity (*p* ≤ 1 × 10^−10^) (Supplementary Fig. 9a-b). For instance, we observed an overlap between strongly associated motifs, e.g., ZIC3, ZNF136, RFX6, and CRX, and nucleotides in the human frame 2 multiple sequence alignment associated with MPRA activity (Supplementary Fig. 9c). We also observed an enrichment of GATA5 and ZNF317 in both MER11_G1/4 which were due to apparent motif turnovers at different locations. When we compared the specificity of enriched motifs, we consistently observed a higher motif enrichment amongst MER11 new subfamilies relative to the originally assigned subfamilies (Fig. 3g and Supplementary Fig. 6h). For example, the highest proportion of frame 2 sequences containing TEAD1 was 66.5% amongst new subfamilies and 40.5% amongst original subfamily annotations, while the lowest proportion was 2.4% amongst new subfamilies and 15.3% amongst original annotations. Taken together, we concluded that the four new subfamilies of MER11 were distinguishable based on distinct sets of motifs associated with MPRA activity.

### The human MER11 new subfamilies are conserved in the primate lineage but spreading independently

To further characterize the evolution of the new subfamilies of MER11, we examined the conservation of instances across the human, chimpanzee, and macaque genomes. We found that MER11 subfamilies had expanded in a lineage-specific fashion since the human-macaque ancestor with, for example, more than 80% of the macaque MER11A sequences absent from the human genome (Fig. 4a). We observed a similar pattern by the alignments of human MER11/34/52 sequences to macaque and vice versa (Supplementary Fig. 10a). In contrast, only 7.4% of the chimpanzee MER11A instances are absent in the human genome (Supplementary Fig. 10b). As expected, the LTR sequences in panTro4 and macFas5 genome builds are conserved among other genome builds (Supplementary Fig. 1c, Supplementary Data 2, Supplementary Data 6 and Supplementary Fig. 10c-d). These species-specific MER11A instances may regulate different sets of genes among primate species (Supplementary Data 7 and Supplementary Fig. 10e). Next, using the approach described above (Supplementary Fig. 3), we built the unrooted tree amongst MER11A macaque instances and identified 33 clusters (11A_m_c1 to c33, Fig. 4b and Supplementary Fig. 10f). Some clusters (e.g., 11A_m_c2/c16/c17/c18) containing > 40% of instances shared with humans were clustered together, while other clusters with a low proportion of instances shared with humans (e.g., 11A_m_c1/c3/c10) were clustered with MER11B/C consensus sequences as labeled in Repbase, suggesting that these instances may be mis-annotated.

**Figure 4.**
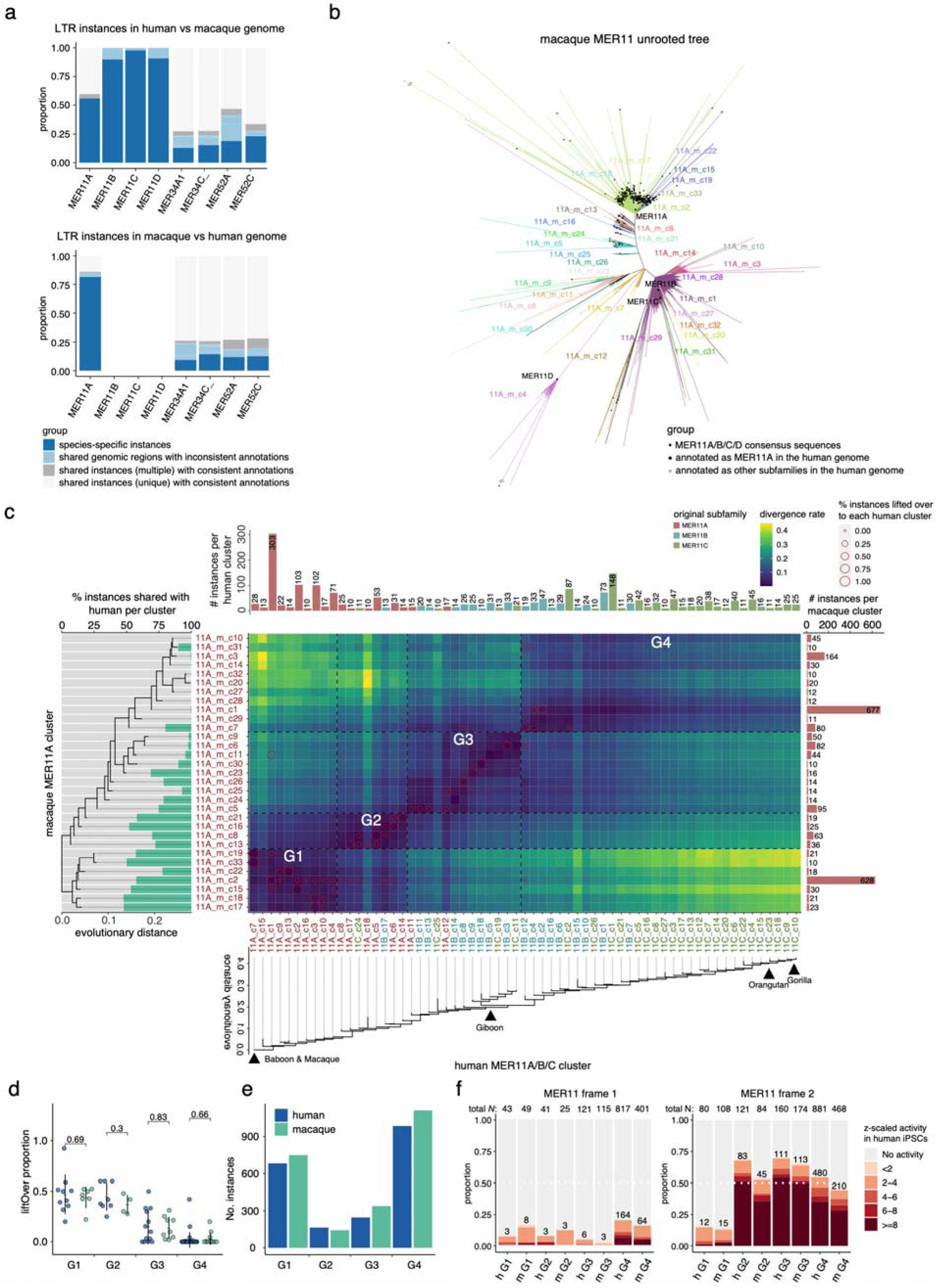
The presence of MER11 new subfamilies in both human and macaque lineages. **a** Conservation and original annotation of MER11 instances in human versus macaque based on liftOver and RepeatMasker. Instances intersected with regions annotated as the same or different subfamilies in another species are shown separately. Instances intersected with unique or multiple regions are also shown separately. **b** Unrooted tree of macaque MER11A instances. MER11A/B/C/D consensus sequences are included as references. Clusters are colored differently. MER11A_m_c4/c12 clusters are clustered with MER11D consensus sequence (Supplementary Fig. 10F) and removed from further analyses. **c** The comparison of MER11 clusters and new subfamilies between humans and macaques. Clusters are arranged by the macaque and human cluster consensus sequence rooted trees. Clusters from every original subfamily are colored differently. The primate lineage when they first integrated is highlighted. Human and macaque new subfamilies are separated by dotted lines. **d** Comparison of the liftOver proportion per new subfamily between human and macaque. **e** Number of instances per new subfamily between human and macaque. **f** Frame1 and frame2 MPRA activity per new subfamily between human and macaque.

Next, we inferred the rooted tree of the macaque MER11A clusters (Fig. 4c, left) and validated that evolutionary old clusters had amongst the highest proportions of instances shared with humans. We found that the MER11 rooted trees had a high consistency between two lineages (Fig. 4c, right). Notably, based on the sequence similarity patterns and the rooted trees, we could also mostly recapture the four new subfamilies we had defined based on the human instances. We further compared the features of each new subfamily between human and macaque lineages. We found that MER11_G1 and MER11_G2 in both species consistently had the highest proportions of instances shared with each other (Fig. 4d and Supplementary Fig. 10g-h). Even though MER11_G4 was the least conserved, we also observed that most instances were in the oldest and youngest groups in both species (Fig. 4e). Finally, we compared the MPRA activities of the frame 1/2 sequences obtained from the two species (Fig. 4f). Overall, the activities of each new subfamily were comparable between two species, with macaque having relatively lower activities (4.4-15.0% for highly active MER11_G2/G3/G4). Taken together, we observed a high conservation in both lineages of the new subfamilies with respect to sequence and MPRA activity. This is especially notable in the youngest new subfamily (G4), in spite of an independent spread following the species divergence.

### The gain of SOX-related motifs within a new subfamily recently occurred in humans and chimpanzees but not in macaques

The gain and loss of TF motifs might help explain the different expansions of MER11 in the primate lineages. To look for such motifs, we focused on MER11_G4 frame 2 sequences in humans, chimpanzees and macaques. Applying the approach described previously, we identified SOX15, POU2F1::SOX2, HSF1, ZBED2, ZNF317, and IKZF1 to be the most associated motifs across three species (Fig. 5a). We then examined the association between the motif combination and MPRA activities amongst MER11_G4 sequences (Fig. 5b). We found that MER11_G4 frame sequences containing either or both POU2F1::SOX2 and SOX15 motifs had the highest activity levels compared to others.

**Figure 5.**
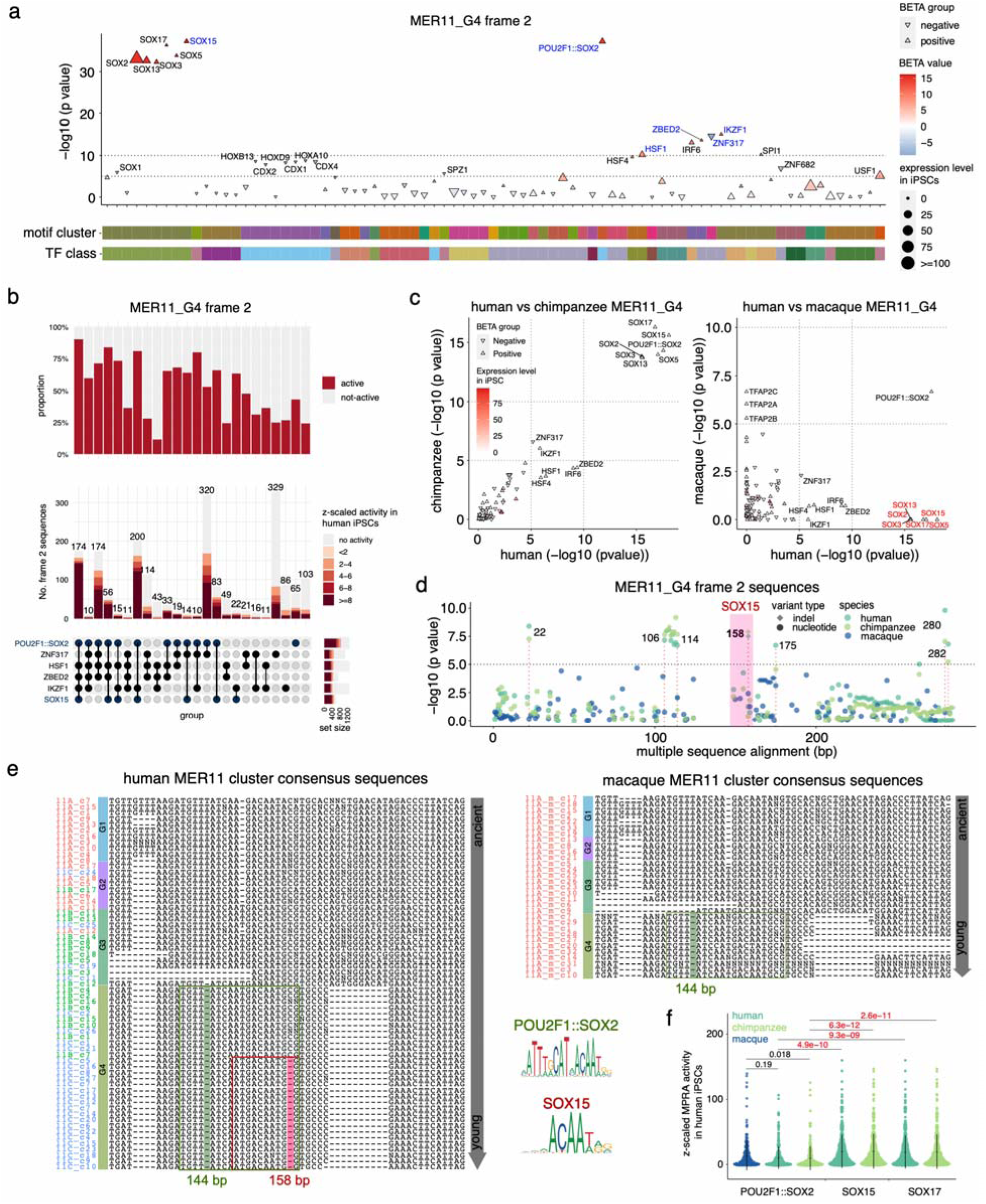
Nucleotide changes and gain of functional motifs during separate expansions of MER11_G4 in primate lineages. **a** Motifs contributing to the activity of MER11_G4 frame 2 sequences. *P* values and effect sizes (BETA values) were computed by the linear regression model using Plink2. **b** Upset plots of different sets of motifs and the proportion of frame 2 sequences with MPRA activity. **c** Association between motifs and the MPRA activity in human, chimpanzee, and macaque MER11_G4 frame 2 sequences. *P* values were computed by the linear regression model using Plink2. **d** Nucleotides associated with the MPRA activity in humans, chimpanzees, and macaques. *P* values were computed by the linear regression model using Plink2. **e** Multiple sequence alignment of human and macaque cluster consensus sequences. Clusters derived from each originally annotated MER11 subfamily are colored differently. New subfamilies are highlighted. POU2F1::SOX2 and SOX15 motifs are highlighted in the boxes. Single-nucleotide deletions associated with the gain of motifs are also highlighted. **f** Comparison of MPRA activity between MER11_G4 frame 2 sequences containing POU2F1::SOX2 and SOX15/17. Sequences containing both motifs are grouped as the sequences with SOX15/17. *P* values were computed using the student’s *t*-test.

Next, we inspected whether the gain of POU::SOX and SOX motifs only occurred in specific primate lineages. We performed the motif association analysis for each species separately. We observed a strong association between the POU2F1::SOX2 motif and MPRA activity consistently in humans, chimpanzees, and macaques; however, SOX related motifs were observed to be significantly associated in humans (*p* ≤ 1 × 10^−10^) and chimpanzees (*p* ≤ 1 × 10^−10^) but were missing in macaques (Fig. 5c). We further analyzed the prevalence of the POU2F1::SOX2 and SOX15/17 motifs amongst the four new subfamilies frame 2 sequences across species. The proportion of sequences containing POU2F1::SOX2 remained below 10% in MER11_G1/2/3 and was significantly increased in MER11_G4, which was consistent across the three species. In contrast, the proportions of SOX15/17 were significantly enriched in humans and chimpanzees but remained low in macaque MER11_G4 (Supplementary Fig. 11a). Only a few other enriched motifs showed a divergence between species with most (e.g., INSM1, DMBX1, and CRX) being consistently enriched (Supplementary Fig. 11b). Nucleotide association further revealed that a human- and chimpanzee-specific single nucleotide deletion at position 158 in the frame alignment was significantly associated with the MPRA activity, which was located within SOX related motifs (Fig. 5d and Supplementary Fig. 11c).

Finally, we inspected the POU::SOX and SOX related motifs across the re-constructed cluster consensus sequences at the target region (Fig. 5e). Human and macaque cluster consensus sequences were compared. We observed a single nucleotide deletion at position 144 occurred between MER11_G3 and MER11_G4 leading to the gain of POU2F1::SOX2 related motifs; another 158-bp deletion occurred between 11B_c7 and 11C_c5 within MER11_G4 leading to the gain of SOX related motifs. Moreover, we compared the MPRA activities between MER11_G4 frame 2 sequences containing POU2F1::SOX2 and SOX15/17 motifs in these species (Fig. 5f). As expected, we observed significantly higher activities in the sequences containing SOX15/SOX17 motifs relative to the sequences containing the POU2F1::SOX2 motif only. Taken together, the phylogenetic analysis of the MER11 family revealed a single nucleotide deletion leading to the gain of SOX-related motifs which was species-specific and significantly increased regulatory potential of the instances in this evolutionary young new subfamily.

### Cryptic endogenous retroviruses subfamilies in the primate lineage with distinct epigenetic profiles

Having shown the usefulness of defining new subfamilies of MER11 to understand the evolution of this family in primate lineages, we wanted to apply the same approach to examine other simian-enriched LTR subfamilies (Fig. 1b and Supplementary Fig. 3 and Supplementary Fig. 4a). Specifically, we first built the unrooted trees and then selected the best representative rooted tree for all analyzed 19 subfamily groups except for the one containing LTR12C and related subfamilies due to practical issues (Methods). In this way, we identified 75 new subfamilies from 18 subfamily groups (containing 53 original LTR subfamilies) (Fig. 6a and Supplementary Data 8). Among them, the LTR7 subfamily group had the most (*N* = 12) new subfamilies and LTR66 remain as a single subfamily. In total, 26 of the individual LTR subfamilies could be subdivided into multiple new subfamilies with a maximum of seven for LTR7C, supporting their high sequence heterogeneity (Fig. 6b). For each LTR subfamily, we selected the new subfamily with the most instances to be the representative for that subfamily. With this, a total of 3,807 (30.0%) instances from these 26 LTR subfamilies were classified into a different new subfamily (Supplementary Fig. 12a). For instance, a total of 258 LTR5_Hs instances (42.7%) were now classified in a non-primary new subfamily.

**Figure 6.**
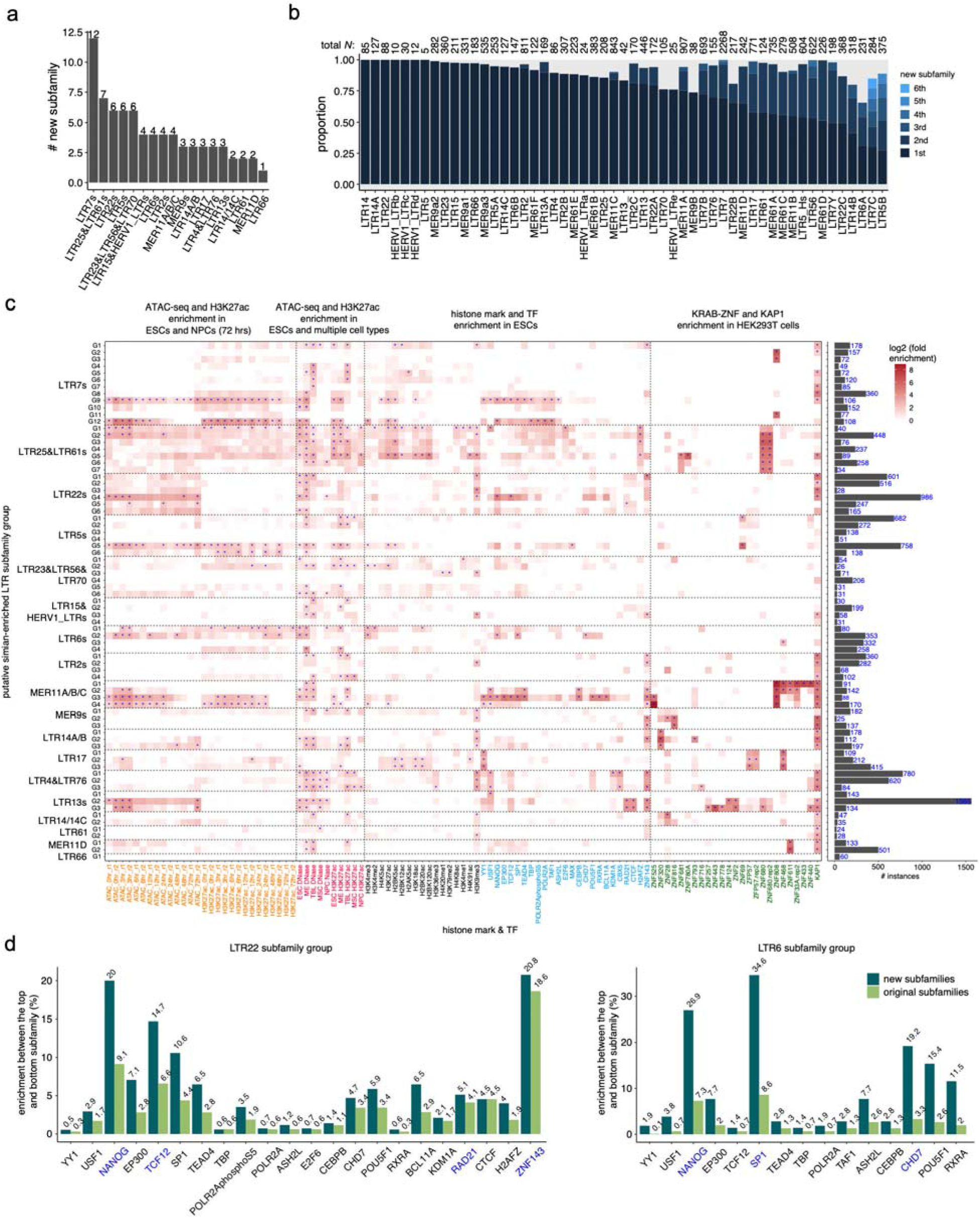
Cryptic endogenous retrovirus subfamilies in the primate lineage with distinct epigenetic profiles **a** Number of newly annotated subfamilies per subfamily group. **b** Proportion of instances per original subfamily classified into the top (representative) new subfamily. The top new subfamily contains the most instances per original subfamily, second new annotation contains the second most instances, and so on. **c** Epigenetic profiles across new subfamilies of 18 simian-enriched LTR subfamily groups. Permutation was used to compute the *p* values. Significantly enriched (log2((actual counts+1)/(mean shuffled counts + 1)) ≥ 1 and *p* value ≤ 0.05) new subfamilies relative to 100 random genomic controls are highlighted. **d** TF specificity of peaks-associated instances between new and original subfamilies in LTR22 and LTR6 subfamily groups. Enrichment was computed as the proportion of peaks-associated instances in the top new subfamily, the one with the highest proportion of peak-associated instances, minus the proportion in the bottom new subfamily, the one with the lowest proportion, for each TF. The same was calculated between the top original subfamily and the bottom original subfamily. Blue color indicates TFs significantly enriched in any new subfamily.

To validate the approach, we reanalyzed the LTR7 subfamily which was carefully studied through phylo-regulatory analysis (Carter et al. 2022). Here, we also included other related subfamilies from the LTR7 subfamily group (i.e., LTR7B, LTR7A, LTR7Y) to depict their full evolutionary history (Supplementary Fig. 12b). As expected, we observed a high consistency between the 12 new subfamilies we identified and the sequence clusters previously reported by Carter et al.(*42*) (Supplementary Fig. 12c). Moreover, by looking at the epigenetic profiles across different cell types, we found that the evolutionary young clusters from LTR7_G12, similar to LTR7up1/up2/up3 from Carter et al.(*42*), were more active and enriched for several TFBSs as compared to other new subfamilies in ESCs (Supplementary Fig. 12c). Additionally, LTR7_G4/G5 were found to be enriched for accessibility and H3K27ac peaks in trophoblast cells (TBL) compared to other cell types, while LTR7_G9 was enriched for accessibility for mesendoderm cells compared to others. We also observed the enrichment of ZNF808 binding sites in LTR7_G3/G11 and KAP1 binding sites in LTR7_G7/G8. These results highlight that the LTR7 new subfamilies have distinct cell-type specific epigenetic profiles.

Finally, we explored the epigenetic properties of the 75 newly annotated subfamilies (Fig. 6c and Supplementary Data 9). As expected, we found that new subfamilies of many subfamily groups were significantly active during the differentiation from human ESC to NPCs, such as LTR5s, LTR6s, LTR7s, LTR13s, LTR22s, and MER11s. LTR new subfamilies were also active in a cell-specific manner, such as LTR6_G4 in mesendoderm cells (ME). We further looked at the epigenetic states in ESCs and found that they were also enriched for different sets of histone marks and TFBSs between new subfamilies. For instance, LTR13_G9/G12 were enriched for RAD21, CTCF, and ZNF143 except for LTR13_G1. We also observed a distinct enrichment of KRAB-ZNF and KAP1 binding sites in HEK293T cells between new subfamilies within certain subfamily groups, such as MER9s. Focusing on the six LTR5 new subfamilies revealed that LTR5_G5 had amongst the highest epigenetic enrichment (Supplementary Fig. 12d). LTR5_G1/G2/G3, which were evolutionarily older, were also overrepresented for multiple active histone marks. As expected, we also observed a higher TF specificity amongst new subfamilies relative to the subfamilies, such as LTR6 and LTR22 subfamily groups (Fig. 6d). Taken together, the newly annotated simian-enriched LTR subfamilies were enriched for different sets of active histone marks, TFs, and KRAB-ZNFs, suggesting differential evolutionary trajectories.

## DISCUSSION

The MER11 family of endogenous retroviruses was previously analyzed for its phylogeny and TF binding sites and motifs(*45*, *46*, *56*, *57*). Consistently, MER11A was found to be more ancient compared to MER11B based on the LTR sequences of 78 human HML8 proviral sequences, and MER11C is the youngest (*45*). They also observed that MER11A/B/C may not be monophyletic groups. However, these studies lacked the assays demonstrating the regulatory activity of distinct genomic variants, nor elucidate the evolutionary process of their regulatory function. Here, based on a critical assessment of the available annotations of MER11 instances, we demonstrate the importance of analyzing the regulatory activity of thousands of genomic instances (copies) using properly reconstituted subfamilies. We found such an annotation critical to understand the expansion process and diverse co-option strategies (e.g., coding and nocoding transcripts, cis-regulatory elements and 3D chromatin organization) of MER11 across diverse primate genomes(*5*, *40*). Moreover, such a phylo-regulatory approach allowed us to detect stronger and new associations between newly annotated MER11 subfamilies and their epigenetic profiles. Following these observations, an MPRA helped us measure the regulatory activity of MER11 variants at an unprecedented scale. MPRAs were previously used to analyze the regulatory activity and evolution of other LTR families, such as LTR18A(*55*), however none of these studies focused on refining the annotation of the instances themselves.

Among the four new subfamilies of MER11, we found that the intermediate-aged MER11_G2/G3 but not MER11_G1 (oldest) contained multiple TF motifs, such as ZIC and TEAD (Supplementary Fig. 9c). These TF motifs were conserved between human and macaque (Supplementary Fig. 11b), suggesting a functional role during the early expansion of MER11, and before the divergence between these two species. It remains unclear whether these motifs were originally present in the ancestral MER11 sequence but subsequently lost in G1, or if they were gained in G2/G3. In contrast, MER11_G4 did not have the ZIC and TEAD motifs, but gained the POU::SOX motif following a single nucleotide deletion. This change might have played a role in the expansion of MER11_G4 in the simian ancestral genome. Notably, in human MER11_G4, another deletion occurred within the same motif region and was detected to be a sole SOX motif in our analysis (Fig. 5e). This suggests an increase in binding affinity for the SOX proteins, supported by the observed increase in the MPRA activity (Fig. 5f). This deletion emerged after the divergence between humans and macaques, potentially contributing to the human-specific expansion of the younger MER11_G4. These results demonstrate that using the reconstituted MER11 subfamilies combined with MPRA can shed light into the process of LTR expansion and divergence at the single nucleotide level. The enhancer function of the MER11 enhancer in relation to the host genome awaits validation in future studies.

The enhancer function of LTRs is influenced not only by transcription activators but also by the de-repression of KRAB zinc-finger transcription repressors that play a role in the defense mechanism of the host genome(*58*, *59*). In our study, we observed that various KRAB repressor binding sequences were differently enriched in the new subfamilies of MER11 (Fig. 2b). This suggests that proper classification and annotation could also be useful to understand the arms race between transcriptional activation and repressive mechanisms. Previous research reported ZNF808 binding to MER11 during pancreatic development(*57*). It remains uncertain which ZNF protein(s) potentially bind to MER11 in iPSCs, although we found a motif for ZNF136, which is expressed in iPSCs, in the frame 2 region of MER11_G2/G3. However, it is important to note that in our MPRA experiment, we primarily focused on the core region of the chromatin accessibility (around 250 bp), which may have limited our ability to fully detect ZNF motifs or mutations that negatively correlate with MPRA activity and contribute to the de-repression. To understand the molecular mechanism of the arms race between transcriptional activation and repressive mechanisms (e.g. mutations in ZNF motifs), a broader range of ERV/LTR sequences should be explored.

Even though the LiftOver method and chain files are commonly used in analyzing transposable elements in primate genomes(*42*, *55*, *60*, *61*), newer assemblies and whole genome alignments could be further used to improve the accuracy in near future. Our approach with MER11 highlights the significance of the phylogenetic classification and annotation of LTRs to understand their functional evolution. Unfortunately, the current annotation of LTR subfamilies groups together heterogeneous sets of sequences (Fig. 1b). We also observed that macaque MER11B/C/D instances were all mis-annotated as MER11A (Fig. 4a-b), which suggests that non-human species may be even more problematic. This issue is likely particularly prevalent in primate LTRs due to their high degree of sequence similarity. To highlight this phenomenon beyond MER11 subfamilies, we used a similar approach across the 53 simian-enriched LTR subfamilies and observed that nearly half (26 out of 53) encompasses new annotations. This analysis suggests a new annotation for 30% of the genomic instances from these 26 simian-enriched LTR subfamilies (Supplementary Data 3). As for MER11, these refined annotations could reveal signals that were missed previously and foster new discoveries relative to the contribution of TEs to primate genome evolution.

Finally, the classification and annotation of TEs across species has been a challenging problem because of the lack of ground truth(*34*). Going forward, we argue that using repeat instances epigenetic and functional profiles, as we have done here, could be an effective strategy to evaluate alternative methods for TE classification and annotation.

## METHODS

### LTR phylogenetic analysis

#### LTR orthology and sequence divergence analysis

To study the expansion of LTR subfamilies in primate lineages, we lifted over the human LTR instances to representative primate species and the mouse. The human (hg19) TE annotation file “hg19.fa.out” was obtained from the UCSC database (http://hgdownload.soe.ucsc.edu/goldenPath/hg19/bigZips/), which also contains the divergence rates (% substitutions) of instances relative to each subfamily consensus sequence. After renaming few subfamilies, we then converted the downloaded “.out” file to BED format using makeTEgtf.pl (https://github.com/mhammell-laboratory/TEtranscripts/issues/83) script. Chain files from human (hg19) to chimpanzee (panTro6), gorilla (gorGor3), orangutan (ponAbe2), gibbon (nomLeu3), macaque (macFas5), baboon (papAnu2), marmoset (calJac3), lemur (micMur1), and mouse (mm10) were downloaded from the UCSC database (http://hgdownload.soe.ucsc.edu/goldenPath/hg19/liftOver/). We also lifted over hg19 to hg38 as a control. BnMapper (https://github.com/bxlab/bx-python) with the parameters “*-k -t 0.5*” was used for the liftOver analysis. Simian-enriched LTR subfamilies were detected as the LTR subfamilies with a minimum of 100 instances (≥ 200 bp) and a maximum of 20% instances that were shared with the lemur genome. To identify subfamilies with potential distinct expansion in primate lineages, we further kept the subfamilies with a maximum of 60% instances that were also present in the macaque genome.

We then explored the orthologous of LTR sequences across more primate genomes. Briefly, 77 assemblies of a total of 47 primate species were downloaded from various databases (Supplementary Data 1). After extracting the human (hg19) LTR sequences together with ± 1 kb adjacent sequences using Bedtools *slop* and *getfasta* functions(*62*), we aligned them against each of the primate genomes using minimap2 with paramameters *“-ax map-pb*”(*63*). We then kept top alignments with MAPQ ≥ 5 and mappable length of ≥ 30% LTR sequences and mappable adjacent sequences (either upstream or downstream) ≥ 300 bp.

Evolutionary ages were computed based on the divergence rates as we previously described(*64*, *65*). Briefly, the divergence rate of each LTR instance relative the corresponding consensus sequence was computed by RepeatMasker. The divergence rates were first divided by the substitution rate for the human genome (2.2×10^−9^) and then averaged across instances from each subfamily as its evolutionary age in million years.

#### LTR consensus sequence similarity and network analysis

We first retrieved the TE consensus sequences in FASTA format from the Repbase database(*66*), which was used to annotate the human (hg19) and macaque (macFas5) genomes used here. We then calculated the sequence similarity score by comparing the sequences amongst themselves using Blastn (BLAST 2.13.0+)(*67*) with the parameters of “*-task dc-megablast -outfmt 6 -num_threads 4*” with the default cut-off (E-value < 10).

After that, the resulting bit scores between each pair of sequences were used as the input of Cytoscape (v3.10.0) for the network analysis(*47*, *68*). Subfamilies with similar consensus sequences were categorized into a subfamily group for the following analysis. Specifically, we first used the edge score of 200 to identify confident subfamily groups containing each candidate simian-enriched LTR subfamily. We then used all edges to recover closely related subfamilies for groups containing a single candidate LTR subfamily. SVA subfamilies are homologous to both LTR5_Hs and Alu sequences in different regions, thus we only kept LTR5 subfamilies within this subfamily group for the following analysis.

#### Human MER11 unrooted trees reconstruction

We first extracted the coordinates of instances (≥ 200 bp) from each MER11 subfamily from the TE annotation BED file separately. We then used Bedtools2 *getfasta* function to extract sequences from the human reference genome (hg19) with the parameter “*-nameOnly -s*”. To reconstruct the evolutionary tree, we first performed the multiple sequence alignment of instances from each subfamily using MAFFT (v7.5.05)(*69*) with the parameters “*--localpair --maxiterate 1000*”. The sequence alignment was further refined using PRANK (v170427)(*70*) with the parameters “*-showanc -njtree -uselogs -prunetree -F -showevents*”. We then used trimAL (v1.4.1)(*71*) to remove gaps that were present in less than 10% of sequences. Lastly, the evolutionary tree was obtained by IQ-TREE 2 (v2.1.2)(*72*) with the parameters “*-nt AUTO -m MFP -bb 6000 -asr -minsup .95 -T 4*”.

#### Human MER11 clusters detection and consensus sequences median-joining network analysis

To determine MER11 clusters per subfamily, we identified branches supported by > 95% ultrafast bootstrap and a minimum internal branch length of 0.02 to any other instances and contained ≥ 10 instances. The original MER11A/B/C/D consensus sequences were also included as references and instances from the identified branches containing < 10 instances were excluded in the following analyses. We then used the *consensusString* function from Biostrings (v2.64.1) R package (https://bioconductor.org/packages/Biostrings) to get the consensus sequences with the majority rule (> 0.51). The cluster consensus sequences were then submitted to PopART (v1.7)(*73*) for the median-joining network analysis. MER11D clusters were also included to confirm their distal relationships with other MER11A/B/C clusters.

#### Human MER11 cluster consensus sequences divergence rate analysis

We used MAFFT with the parameters “*--globalpair –maxiterate 1000*” to align MER11 cluster consensus sequences with the default parameters. We then used RAxML (v8.2.12) with the parameters “*raxmlHPC-PTHREADS-AVX -f x -p 12345 -m GTRGAMMA*” to compute the divergence rates (maximum likelihood distances) between each pair of human cluster consensus sequences. We also re-calculated the divergence rates of every instance versus their corresponding original and new subfamily consensus sequences separately. We used the consensus sequence of relatively ancient cluster (i.e., 11A_c7, 11A_c8, 11B_c11, and 11C_c2) to represent each new subfamily. We first ran RepeatMasker with all MER11A/B/C instances and each consensus sequence as the inputs and parameters “-e rmblast -pa 4 -s -no_is”. We then used the “one_code_to_find_them_all_but_sanely.pl” (https://github.com/mptrsen/mobilome/blob/master/code/Onecodetofindthemall/) script to combine adjacent partial hits. We used ggplot2 for the visualization.

#### Human MER11 new subfamilies determination

We next want to infer the best rooted tree among all MER11A/B/C cluster consensus sequences. Firstly, we performed the multiple sequence alignment analysis using MAFFT with the parameters “*--localpair --maxiterate 1000*”. Secondly, the alignment was refined using PRANK with the parameters “*-showanc -njtree -uselogs -prunetree -F -showevents*”. After that, we constructed the rooted trees using IQ-TREE 2 with the parameters “*--model-joint 12.12 -B 1000 - T AUTO --root-test -zb 1000 -au*”. It also performed the statistical tests for rooting positions on every branch. We then selected the best tree based on the ranking, different statistical tests, and the liftOver rates to other primate species. Specifically, we selected from the top-ranked rooted tree, which rooted cluster has among the highest liftOver rates in the most ancient primate species we used above. We also prioritized the rooted trees having the highest values amongst different statistical tests. The top-selected rooted tree was then used in the following analyses. The branch length (bootstrap value) from each cluster to the root referred to the evolutionary age.

We next determined the new subfamilies based on the internal branch lengths of the top-selected rooted tree. Since the branch lengths varied between subfamily groups, we grouped clusters as a new subfamily which have amongst the top branch lengths to others. We also confirmed the new subfamilies by examining the pair-wide divergence rates to look at extreme values between every adjacent clusters. We lastly kept the new annotations containing a minimum of 25 instances.

#### Macaque MER11A phylogenetic analysis

We performed the same phylogenetic analysis for macaque (macFas5) MER11A instances (≥ 200 bp). After we subdivided them into clusters, we also inferred the cluster consensus sequences rooted tree using the same approach. We then determined new subfamilies for the originally annotated macaque MER11A instances while two new subfamilies were named “G4-1” and “G4-2” according to their close relationship with human “MER11_G4” consensus sequences. We also lifted over the instances from each cluster to human (hg19) using bnMapper with the parameters “*-k -t 0.5*”. The chain file “macFas5ToHg19” was downloaded from the UCSC database (https://hgdownload.soe.ucsc.edu/gbdb/macFas5/liftOver/). The divergence rates between human and macaque MER11 cluster consensus sequences were also computed using the same approach above. We used ggplot2 for the visualization.

#### Simian-enriched LTR subfamily groups phylogenetic analysis

We performed a similar analysis for other simian-enriched LTR subfamily groups. Briefly, we first constructed the unrooted trees of instances (≥ 200 bps) for every subfamily from a group. After identifying the clusters per subfamily, we constructed the rooted trees and selected the best one to determine new subfamilies for each subfamily group. Using the same approach (Supplementary Fig. 3), we kept new subfamilies with a minimum of 25 instances except LTR61 subfamily which has a small number of instances.

### LTR epigenetic analysis

#### Chromatin accessibility permutation analysis at the TE subfamily level

We downloaded the ATAC-seq peaks from the NCBI Gene Expression Omnibus (GEO) database (GSE115046)(*51*). After that, we used the same approach as we previously described to evaluate the enrichment level relative to the random genomic background per TE subfamily(*65*). Briefly, we first shuffled the peak regions 1000 times relative to the distribution of peaks. We then computed the number of instances per subfamily that overlapped with the actual and shuffled peak summits separately using Bedtools2(*62*) *intersect* function with the parameters “*intersect -wa -u -a*”. After that, we counted the number of instances that were associated with peaks per new subfamily. Fold enrichment was computed as the number of actual peaks-associated instances divided by the average number of shuffled peaks-associated instances, and the permutation test was used to compute the *p* values. We also performed the same analyses on another H1 ESC (E003) DNase-seq dataset(*52*).

#### Differential accessibility and H3K27ac activity between ESCs and NPCs

To identify LTR subfamilies with the accessibility and H3K27ac activity changes during the cell differentiation, we re-analyzed the accessibility and H3K27ac activity changes in TEs between ESCs and NPCs. The additional ATAC-seq and H3K27ac peak files were downloaded from the same sources. Fold change per subfamily between ESCs and NPCs was computed as we described previously(*65*). We then kept subfamilies with a minimum of two-fold enrichment of chromatin accessibility in ESCs compared to NPCs. We further validated the results in other differentiated cells including mesendoderm cells (ME, E004), trophoblast-like cells (TBL, E005), mesenchymal stem cells (MSC, E006), and neural progenitor cells (NPC, E007)(*52*). We kept the LTR subfamilies that were significantly enriched in the accessibility and H3K27ac in ESCs from two independent sources.

#### Permutation test of other epigenetic marks overlapped each MER11 subfamily

Except the above datasets, we further obtained additional H1 ESC histone and TF Chip-seq peaks (https://www.ncbi.nlm.nih.gov/geo/roadmap/epigenomics/), and HEK293T KRAB-ZNF and KAP1 Chip-seq peaks(*56*). We then examined the enrichment of each epigenetic mark overlapped with MER11 subfamilies using the same approach. The number of peaks overlapped with each subfamily was normalized by the total number of peaks per mark. We then kept epigenetic marks overlapped with a minimum of 20 instances per subfamily that were significantly enriched (fold enrichment ≥ 2 and *p* value ≤ 0.05).

#### Hypergeometric test of histone marks and TFBSs at the cluster level

We selected the significantly enriched epigenetic marks in any MER11 subfamilies as well as other well-known active (H3K4me1/2) and repressive marks (H3K27me3). We then inspected whether these marks were overrepresented within specific MER11 clusters. To do it, we intersected the actual peaks with MER11 clusters using Bedtools2 *intersect* function with parameters “*-wa -e -f 0.5 -F 0.5 -u*”. After that, we computed the proportion of instances per cluster were overlapped with each peak. Moreover, we used the hypergeometric test (R *phyper* function) to compute the *p* value for the enrichment of peaks-associated instances per cluster relative to each subfamily. The *p* values were adjusted using R *p.adjust* function with the *Benjamini & Hochberg* method.

#### Permutation test of epigenetic marks overlapped with new subfamilies of each subfamily group

We examined the enrichment of the above epigenetic marks overlapped with each determined MER11 new subfamily. Specifically, we first shuffled the peak regions 100 times relative to the distribution of peaks using our previous approach(*65*). We then intersected the actual and shuffled peaks with instances from each new subfamily using Bedtools2 *intersect* function with parameters “*-wa -e -f 0.5 -F 0.5 -u*”. After that, we counted the number of peaks-associated instances per new subfamily. Fold enrichment was computed as the number of actual peaks-associated instances divided by the average number of shuffled peaks-associated instances. A permutation test was used to compute the *p* values. New subfamilies with a minimum of two-fold enrichment (log2((actual counts+1)/(mean shuffled counts + 1)) ≥ 1) with *p* value ≤ 0.05 were kept. We also filtered out epigenetic marks overlapped with less than five instances from new annotations containing a maximum of 100 instances and less than 5% for new subfamilies containing more than 100 instances separately.

### LTR lentiMPRA library design

#### Detection of sequence frames along the LTR consensus sequences

The computed reads per million (RPM) distribution of accessible MER11B instances were first aligned to each subfamily consensus sequence using the downloaded “.align” file as we previously described(*65*). We determined the consensus accessible regions based on the aggregated RPM distribution along the consensus sequences. After that, we extracted the consensus accessible sequences (frame regions) centered at the peak summits at around 250 bp long. We also retrieved the homologous MER11B and MER11C consensus sequences based on the multiple sequence alignment using ClustalW2 with the default parameters (https://www.ebi.ac.uk/Tools/msa/clustalw2/). Similarly, we determined the frame regions on MER34, and MER52 subfamilies separately.

#### Extraction of human, chimpanzee, and macaque LTR genomic sequences

We used the in-house Python script “*Organize_seqFile_to_consensus.py*” to retrieve annotated MER11, MER34, and MER52 sequences that were homologous to each frame sequence. We then kept homologous sequences with a maximum of 270 bp and without ambiguous nucleotides “N”. Sequences that were shorter than 70% of the frame sequences were removed. Sequences with a reverse alignment against the frame sequences were converted to the reverse complementary sequences. Similarly, we obtained the chimpanzee and macaque genomic sequences homologous to each frame sequence. Chimpanzee (panTro4) and macaque (macFas5) TE annotation files (Repeat library 20140131) were downloaded from http://www.repeatmasker.org/species/macFas.html and http://www.repeatmasker.org/species/panTro.html.

#### Negative and positive controls

We first used the shuffleFasta tool with the parameter “*-n 100*” to randomly select 100 sequences from the extracted genomic sequences (https://krishna.gs.washington.edu/content/members/vagar/forTaka/). We then used the Python script “*kMerFilter_fromMartin.py*” also from the same link with the parameters “*-k 8 --inclMinOverlap*” to further shuffle the nucleotides. The obtained sequences were used as the negative controls without activities. Sequences with activity(*51*) were also used as positive controls in this study.

#### Consensus sequences

MER11/34/52 subfamily consensus sequences were also included in the library. We also reconstructed the consensus sequences amongst the retrieved genomic sequences per species based on the “.align” file. Consensus sequences with every nucleotide at ambiguous nucleotides “N” that were different from the above genomic sequences were kept.

#### Adding adapter sequence

After the removal of redundant sequences, we added the upstream and downstream adapter sequences “AGGACCGGATCAACT” and “CATTGCGTGAACCGA” to the examined sequences using an in-house Python script.

### lentiMPRA experiment

#### lentiMPRA library cloning and sequence-barcode association

Designed sequence oligos were synthesized by Twist Bioscience. The lentiMPRA library construction was performed as previously described with modifications(*74*). In brief, the synthesized oligo pool was amplified by 7-cycle PCR using forward primer (5BC-AG-f01, Supplementary Data 10) and reverse primer (5BC-AG-r01, Supplementary Data 10) that added mP and spacer sequences downstream of the sequence. The amplified fragments were purified with 1.8x AMPure XP (Beckman coulter), and proceeded to the second round 9-cycle PCR using forward primer (5BC-AG-f02) and reverse primer (5BC-AG-r02, Supplementary Data 10) to add 15-nt random sequence that serves as a barcode. The amplified fragments were then inserted into SbfI/AgeI site of the pLS-SceI vector (Addgene, 137725) using NEBuilder HiFi DNA Assembly mix (NEB, E2621L), followed by transformation into 10beta competent cells (NEB, C3020) using the Gemini X2 machine (BTX). Colonies were allowed to grow overnight on Carbenicillin plates and midiprepped (Qiagen, 12945). We collected approximately 1.2 million colonies, so that on average 70 barcodes were associated with each sequence. To determine the sequences of the random barcodes and their association with each sequence, the sequence-mP-barcode region was amplified from the plasmid library using primers that contain flowcell adapters (P7-pLSmP-ass-gfp and P5-pLSmP-ass-i#, Supplementary Data 10). The PCR fragment was then sequenced with a NextSeq mid-output 300-cycle kit using custom primers (pLSmP-ass-seq-R1, pLSmP-ass-seq-ind1 (index read), pLSmP-ass-seq-R2, Supplementary Data 10).

#### Cell culture, lentiviral infection and barcode sequencing

WTC11 human iPSCs (Coriell Institute, RRID:CVCL_Y803) were cultured on matrigel (Corning, 354277) in mTeSR plus medium (Stemcell technologies, 100-0276) and passaged using ReLeSR (Stemcell technologies, 100-0484), according to manufacturer’s instruction. WTC11 cells were used for the MPRA experiments at passage 43. Lentivirus packaging was performed as previously described with modifications(*74*). Briefly, 50,000 cells/cm2 293T cells (ATCC, CRL-3216) were seeded in four T175 flasks and cultured for 48 hours. The cells were co-transfected with 7.5 μg/flask of plasmid libraries, 2.5 μg/flask of pMD2.G (Addgene 12259) and 5 μg/flask of psPAX2 (Addgene 12260) using EndoFectin Lenti transfection reagent (GeneCopoeia, EF002) according to manufacturer’s instruction. After 8 hours, cell culture media was refreshed and ViralBoost reagent (Alstem, VB100) was added. The transfected cells were cultured for 2 days and lentivirus was filtered through a 0.45um PES filter system (Thermo Scientific, 165-0045) and concentrated by Lenti-X concentrator (Takara Bio, 631232) according to manufacturer’s protocol. We obtained in total 1.2 mL lentivirus solution from the four T175 flasks (100x concentration).

For lentiviral infection, approximately 4 million WTC11 cells per replicate were seeded in mTeSR plus medium supplemented with Y-27632 (Cayman, 10005583) in a 10 cm dish. After 24 hours, the cell culture medium was replaced by fresh mTeSR plus medium without Y-27632. To perform magnetofection, 100 μL/dish of the concentrated lentivirus library, 150 μL/dish ViroMag R/L reagent (OZ Biosciences, RL41000), and 750 μL/dish mTeSR plus medium were mixed and incubated at room temperature for 20 minutes. WTC11 cells in a 10 cm dish were added with the virus-ViroMag mixture and placed on a magnetic plate for 30 minutes. The cells were removed from the magnetic plate, and incubated for 24 hours. The infected cells were cultured for additional 3 days with a daily change of the mTeSR plus media, or induced into neural lineage for 3 days with dual-Smad inhibitors as described previously(*51*). For each experiment, three independent infections were performed to obtain three biological replicates.

DNA/RNA extraction and barcode sequencing were all performed as previously described(*74*). Briefly, genomic DNA and total RNA were purified from the infected cells using an AllPrep DNA/RNA mini kit (Qiagen, 80204). 120 μg RNA was treated with Turbo DNase (Thermo Fisher Scientific, AM1907) to remove contaminating DNA, and reverse-transcribed with SuperScript II (Invitrogen, 18064022) using a barcode-specific primer (P7-pLSmp-assUMI-gfp, Supplementary Data 10), which has a 16-bp unique molecular identifier (UMI). The cDNA and 48 μg genomic DNA from each sample were used for 3-cycle PCR with specific primers (P7-pLSmp-assUMI-gfp and P5-pLSmP-5bc-i#, Supplementary Data 10) to add sample index and UMI. A second round PCR (21 and 26 cycles for DNA and RNA barcode samples, respectively) was performed using P5 and P7 primers (P5, P7, Supplementary Data 10). The PCR fragments were purified and sequenced with a NextSeq high-output 75-cycle kit (15-cycle paired-end reads, 16-cycle index read1 for UMI, and 10-cycle index read2 for sample index), using custom primers (pLSmP-ass-seq-ind1, pLSmP-bc-seq, pLSmP-UMI-seq, pLSmP-5bc-seqR2, Supplementary Data 10).

### MPRA activity measurement

#### Association analysis

To ensure the accurate association between barcodes and LTR inserts with variable lengths, we revised the Python script “*nf_ori_map_barcodes.py*” implemented in MPRAflow (v2.3.1)(*74*). Specifically, we kept reads that were fully matched to the designed oligos nucleotide sequences with the cigar “M” at the extract insert lengths. We then ran the optimized MPRAflow pipeline with the following command and parameters “*nextflow run association.nf –mapq 5 –min-frac 0.5*” to associate each LTR insert sequence with multiple barcodes. Paired-end insert DNA FASTQ files, barcode FASTQ files, and designed library FASTA file were used as the inputs.

#### DNA/RNA count analysis

After that, we ran the MPRAflow command “*nextflow run count.nf –bc-length 15 –umi-length 15 -thresh 5 –merge_intersect FALSE*” to achieve the normalized number of DNA and RNA reads per barcode, and the RNA/DNA ratio associated with each LTR insert sequence. The designed library FASTA, the output file from the above association analysis, and the list of DNA/RNA barcode and UMI FASTQ files were used as the inputs.

#### MPRA activity (alpha value) calculation

We used the MPRAnalyze R package (v1.18.0) to compute the MPRA activity. The DNA and RNA count matrices of three replicates were used as the inputs. We first estimated the library size correction factors (i.e., batch and condition factors) using the *estimateDepthFactors* function with the parameters “*which.lib = “both”, depth.estimator = “uq”*”. After the quant model fitting using the *analyzeQuantification* function, the MPRA activity (alpha value) was obtained using the *getAlpha* function with the parameters “*by.factor = “batch“*”. We then kept LTR insert sequences that were associated with ≥ 10 barcodes in more than two DNA libraries.

#### MPRA activity normalization

We normalized the activity (alpha value) by the negative controls following the same approach here(*51*). Briefly, we first computed the mean absolute deviation of negative control sequences. After we extracted the high quality negative control sequences, we then computed the *z*-scaled MPRA activity using the same formula we previously reported. *P* values were computed based on the standard normal distribution and were further adjusted using the Benjamini–Hochberg method. LTR insert sequences with adjusted *p* value ≤ 0.05 were classified as sequences with MPRA activity.

### Motif and nucleotide association analyses

#### De novo motif discovery and summary analysis

We searched each MER11 frame sequence for known motifs from the JASPAR 2022 database using MEME (v5.2.0) *fimo* function(*75*). After the removal of redundant motifs per frame sequence, we computed the proportion of sequences containing each motif at the cluster level. Similarly, we computed the proportion for each MER11 new subfamily. Human, chimpanzee, and macaque sequences were analyzed separately.

#### Motif association analysis

We implemented a new TE motif association approach to identify motifs that contributed to the activity. The non-redundant motifs per frame sequence were converted to pseudo genotypes for each motif (i.e., presence denoted as A/B and absence denoted as A/A) in “.ped” format. Each row referred to a frame sequence and every two columns (starting from the 7th column) referred to the two pseudo alleles of a motif. The z-scaled MPRA activity was used as the quantitative phenotypes (6th column of the “.ped” file). We also prepared a corresponding “.map” file containing the list of motifs (same order as the “.ped” file). We lastly used Plink2 for the association analysis (https://www.cog-genomics.org/plink/2.0/)(*76*). Specifically, we filtered the genotypes with the parameters “*--maf 0.05 -- make-bed --input-missing-phenotype 999*”. After we computed the allele frequency, the generalized linear model was used for the association analysis with the parameter “*--glm allow-no-covars*”. *P* values and BETA values were visualized by ggplot2. Analyses were done amongst the frame sequences from three species (human, chimpanzee, and macaque) or each species separately.

#### Nucleotide association analysis

We performed the multiple sequence alignment of MER11 frame 2 sequences using MAFFT with the parameters “*--localpair --maxiterate 1000*”. We then converted the variants across each frame sequence into pseudo-genotypes (i.e., minor variant as A/B and major variant as A/A) for each nucleotide along the alignment. Gaps (or indels) and nucleotide changes were analyzed separately. We then used them as the inputs for the association analysis with the *z*-scaled MPRA activity in iPSCs as the quantitative phenotypes. Plink2 was used for the association analysis as we described above. *P* values and BETA values were visualized by ggplot2. Analyses were also done with the inclusion of MER11_G4 frame 2 sequences retrieved from three species or each species, separately.

## ACKNOWLEDGMENTS

We would like to thank Drs. Gracie Gordon and Tal Ashuach for their help with the use of MPRAflow and MPRAnalyze tools. We also acknowledge Institute for the Advanced Study of Human Biology (ASHBi), Institutional Center for Shared Technologies and Facilities of Shanghai Institute of Immunity and Infection, CAS, Calcul Québec and the Digital Research Alliance of Canada for access to computing resources. We thank the Single-Cell Genome Information Analysis Core (SignAC) at WPI-ASHBi, Kyoto University, for their support. We thank Dr. Spyros Goulas for critical reading and suggestions on the manuscript.

## Funding

This work was supported by the World Premier International Research Center Initiative (WPI), MEXT, Japan, the JSPS KAKENHI under Grant Numbers JP21K06119 (F.I.), JP21K15066 (X. C.), the Takeda Science Foundation, Bioscience Research Grants (F.I.), and by the Mitsubishi Foundation, Research Grants in the Natural Sciences (F.I.). This work was also supported by a Canadian Institute of Health Research (CIHR) program grant (CEE-151618). G.B. is supported by a Canada Research Chair Tier 1 award, an FRQ-S, and a Distinguished Research Scholar award. This research was enabled in part by support provided by Calcul Quebec and the Digital Research Alliance of Canada.

## Author contributions

Conceptualization: X.C., G.B., and F.I. Methodology: X.C., Z.Z., G.B., and F.I. Formal Analysis: X.C. and Z.Z. Investigation: X.C., Z.Z., Y.Y., and F.I. Resources: G.B. and F.I. Visualization: X.C. Writing – original draft: X.C. G.B., and F.I. Writing – review & editing: X.C., Z.Z., C.G. G.B., and F.I. Funding Acquisition: X.C., G.B., and F.I. Overall supervision: G.B.

## Competing Interests

F.I. receives funding from Relation Therapeutics.

## Data and materials availability

The datasets generated in this study are available at the NCBI Gene Expression Omnibus (GEO) as accession number GEO: GSE245662. Scripts for main analyses are available at https://github.com/xunchen85/TE-MPRA-and-phylogenetic-analysis and will also be submitted to Zenodo at DOI:10.5281/zenodo.10016500 upon acceptance.

## Notes

### Summary of Updates

Result section 1 was updated; Author affiliations updated; Figure 1 updated; Methods section updated; Supplemental files updated

