## Supplementary Figures for "Cryptic endogenous retrovirus subfamilies in the primate lineage"

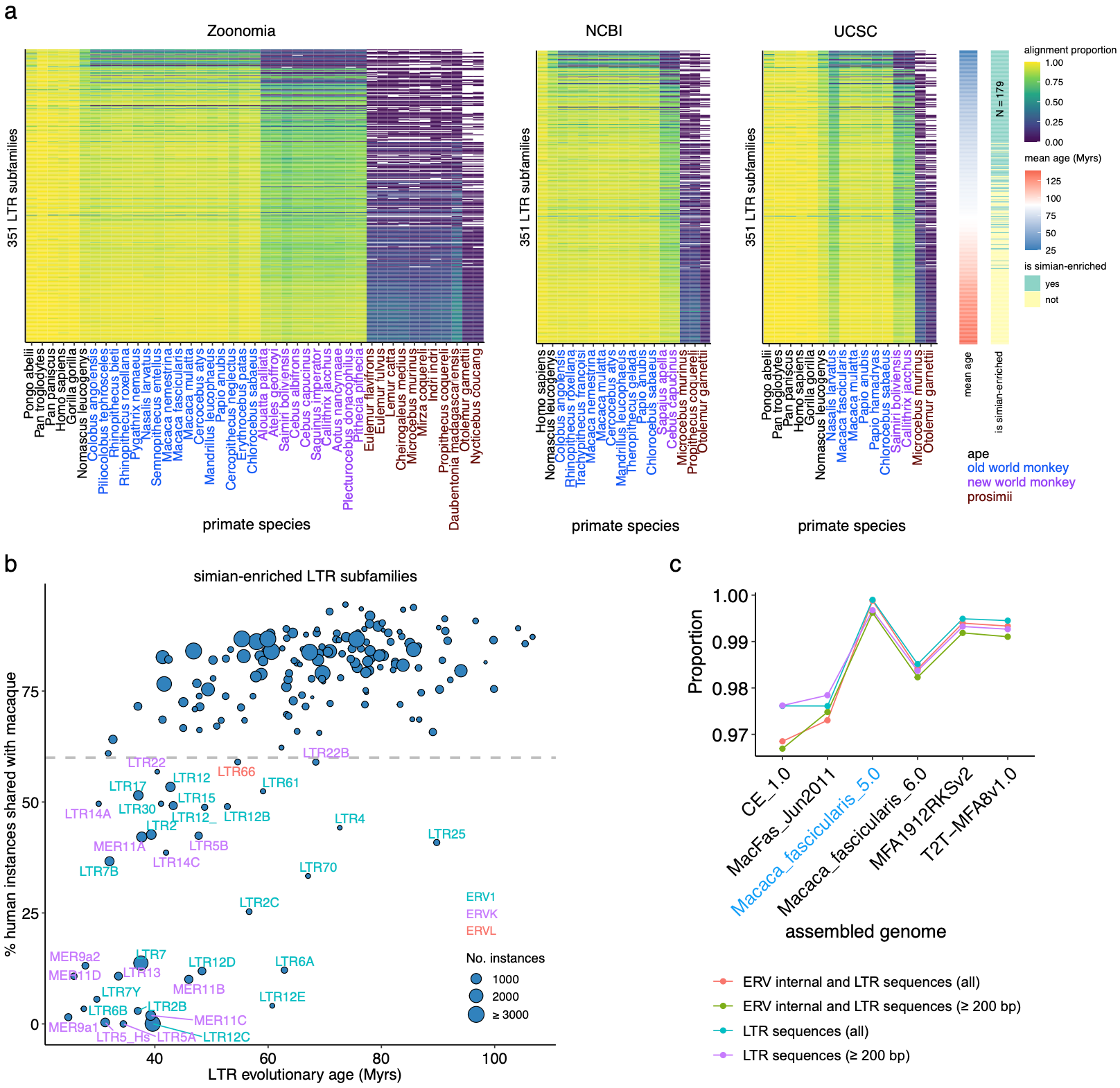

#### Supplementary Figure 1 Detection of evolutionary young simian-enriched LTR subfamilies. a Alignment of LTR subfamilies from human (hg19) to other primate genomes (see Methods). Subfamilies in Fig. 1a were examined. LTR sequences with their adjacent sequences (±1 kb) were used for the alignment by minimap2. b Proportion of instances in the human genome from the 179 simian-enriched LTR subfamilies that are shared with macaques. LTR subfamilies with less than 60% of instances shared with macaques are highlighted. Subfamilies derived from each ERV superfamily are colored differently. c The conservation of annotated macFas5 LTR sequences in other Macaca Fascicularis genome builds. LTR/ERV subfamilies (with or without internal sequences) were calculated separately.

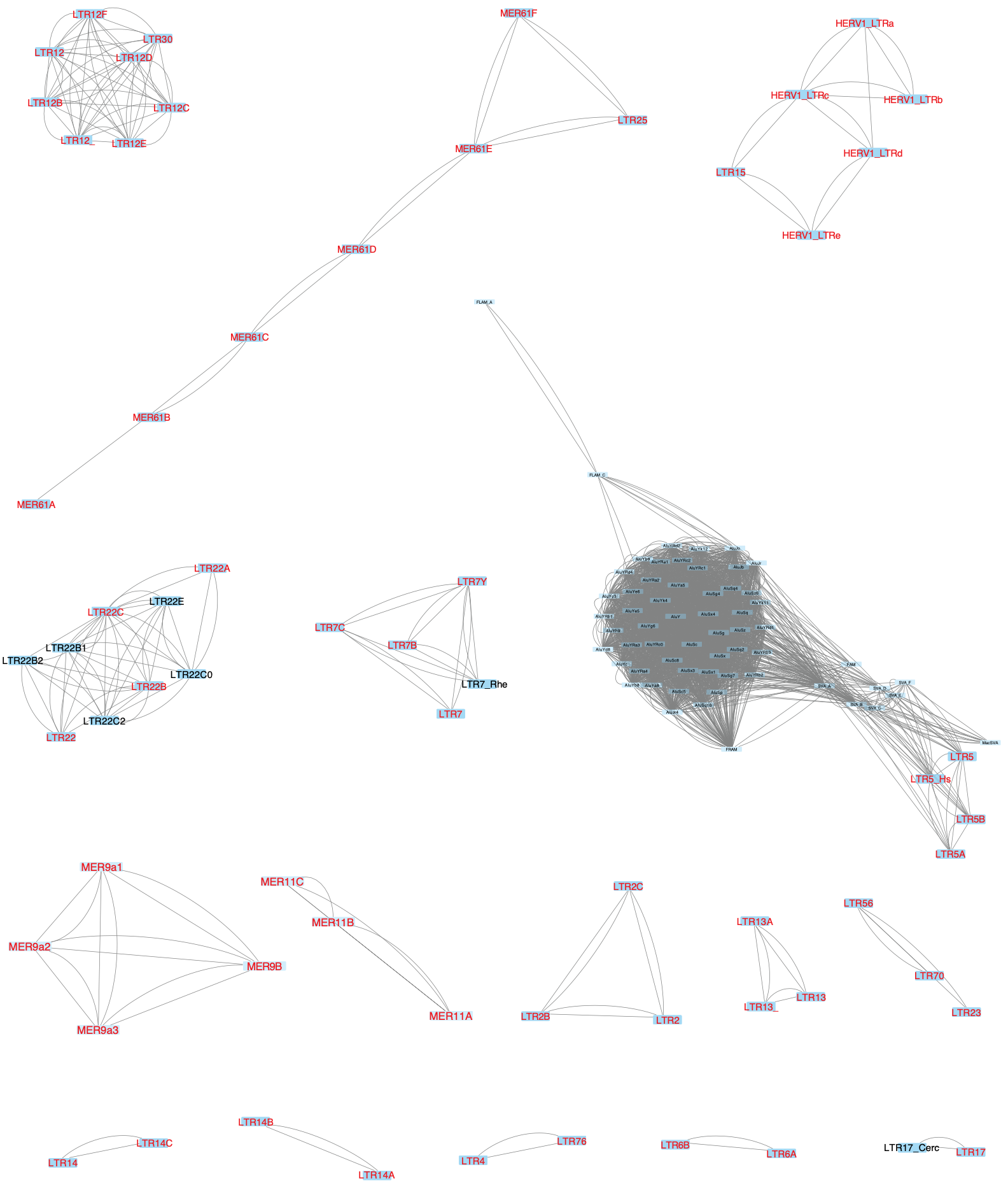

#### Supplementary Figure 2 Putative simian-enriched subfamily groups. Visualization of 16 LTR subfamily groups except three groups with a single subfamily (LTR61, LTR66, and MER11D). Network analysis based on subfamily consensus sequence similarity was used to create clusters (see Methods). Subfamilies that are present in the human genome are highlighted in red. For the subfamily group containing LTR5 subfamilies, Alu and SVA non-LTR subfamilies were excluded from downstream analyses.

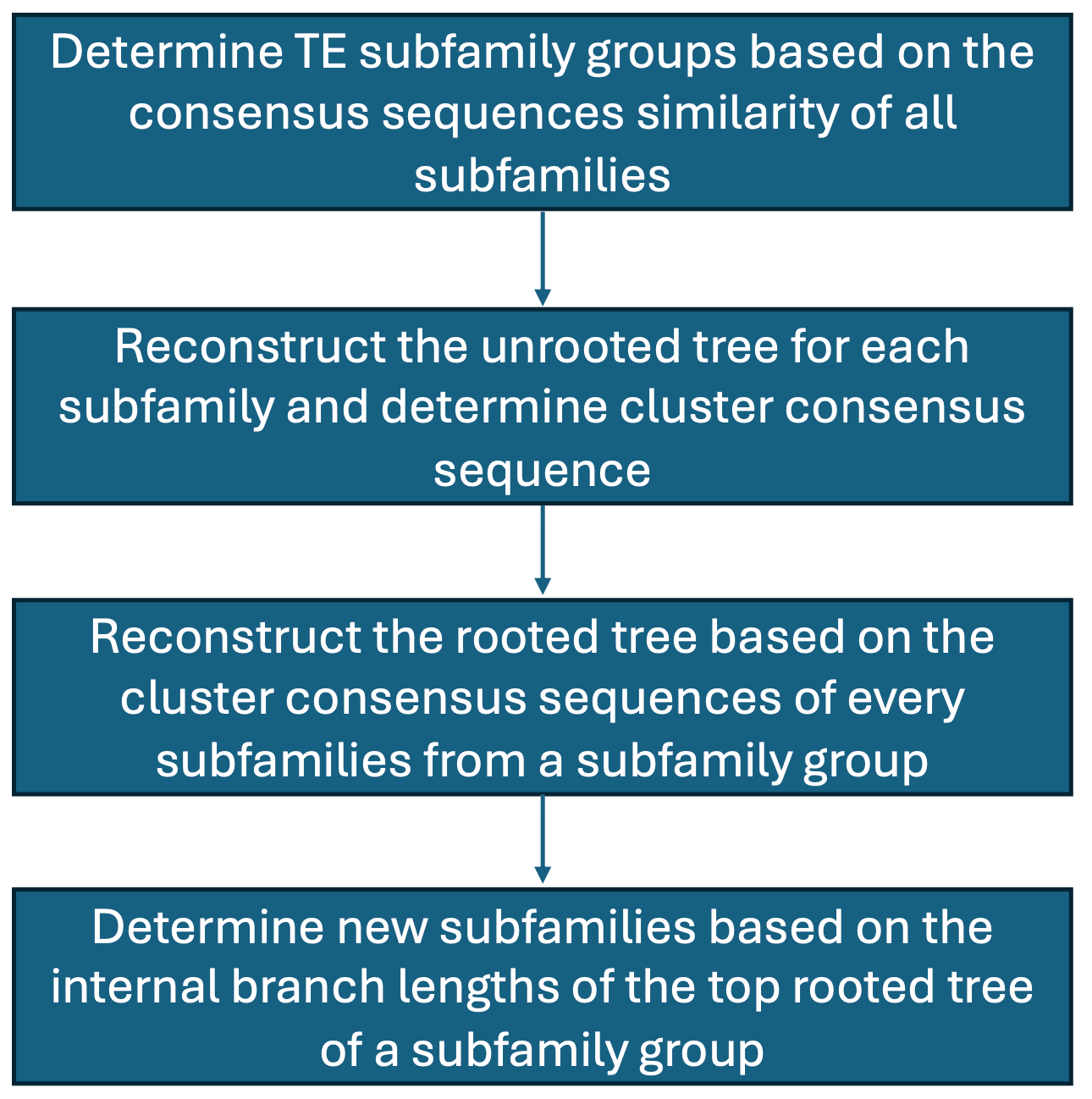

**Supplementary Figure 3** Workflow to classify and annotate the transposable element sequences.

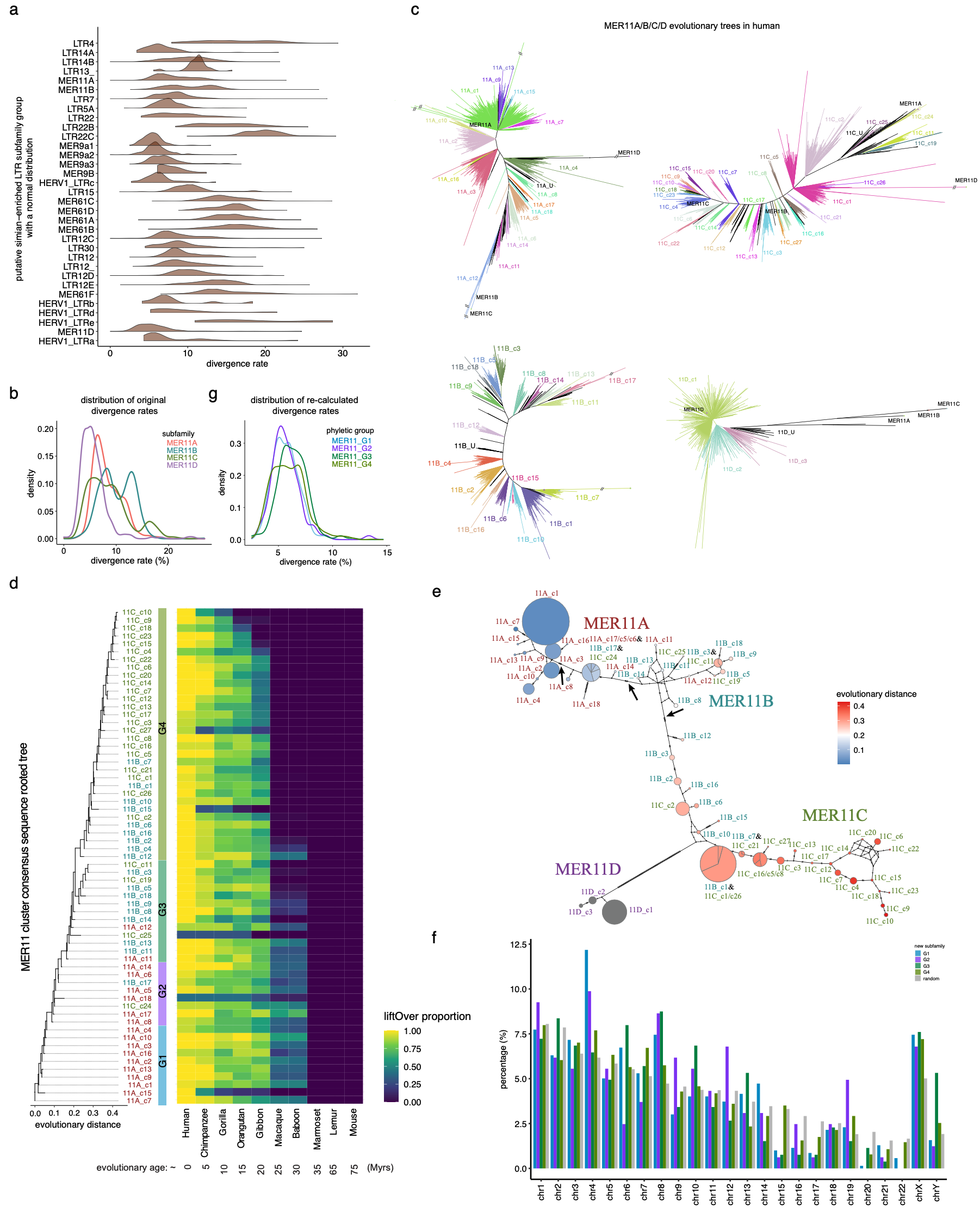

#### Supplementary Figure 4 Evolution of human MER11 subfamilies. a Divergence rate distribution of instances relative to the subfamily consensus sequence. Subfamilies that have expected distributions (Chi-square test, Bonferroni adjusted *p* values ≥ 0.001) are shown. b Divergence rate distribution of MER11 subfamilies. c Unrooted trees of MER11A, MER11B, MER11C, and MER11D subfamilies. Human instances ≥ 200 bp were analyzed. Consensus sequences are in black and newly identified sequence clusters are in different colors. d Selected rooted tree containing 63 MER11A/B/C cluster consensus sequences and the proportion of instances shared with mouse and other primate lineages. e Median-joining network of 66 MER11 cluster consensus sequences. Color refers to the evolutionary age. Size refers to the relative number of instances amongst clusters. The arrow indicates the edges have the most mutations between new subfamilies. f Distribution of MER11_G1/G2/G3/G4 instances across the human genome. Random refers to the expected value which was computed by dividing the length of each chromosome by the cumulative length of all chromosomes. g Distribution of re-computed divergence rates of instances relative to each new subfamily consensus sequence (see Methods).

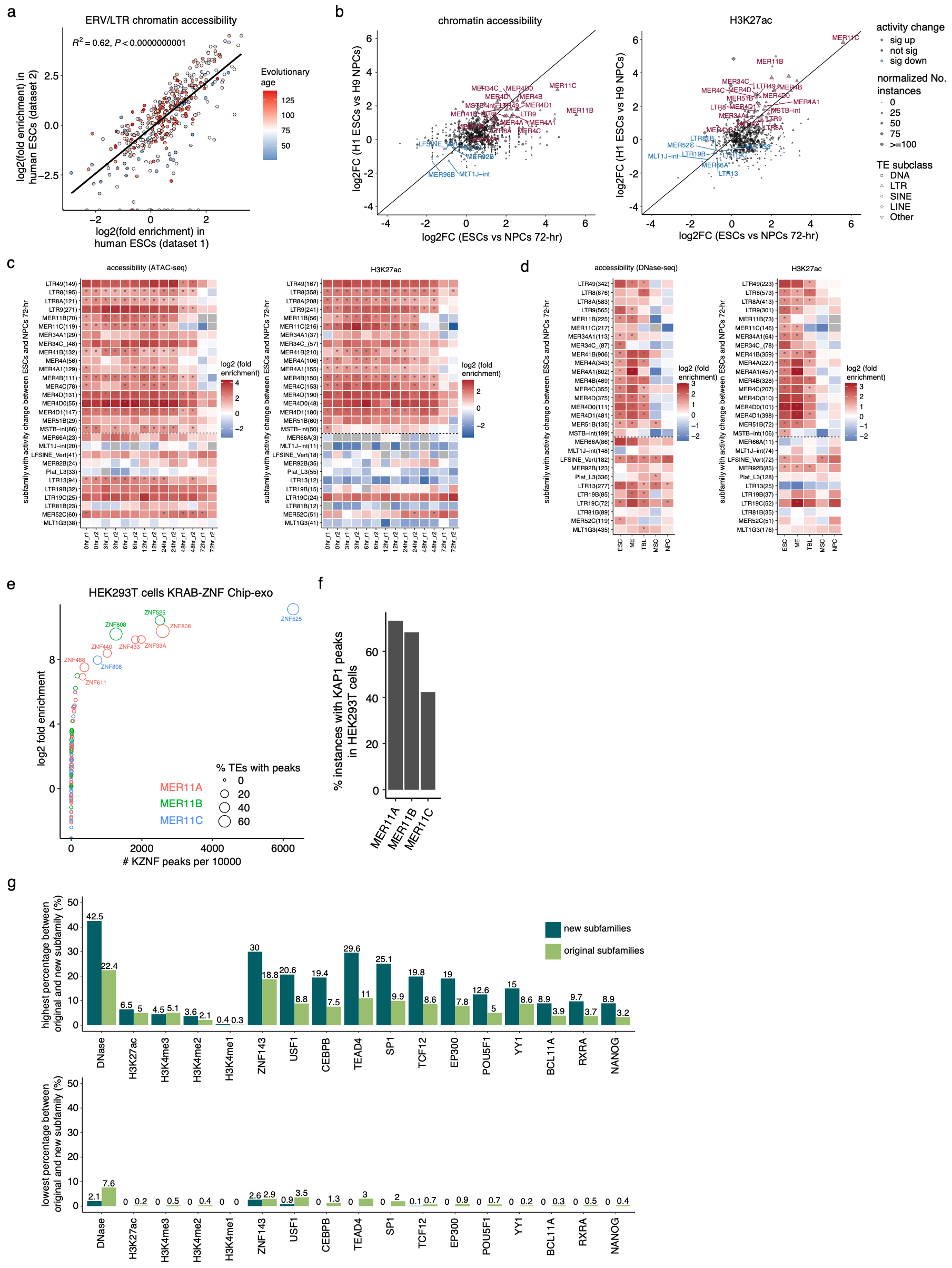

#### Supplementary Figure 5 Epigenetic profiles of human MER11A/B/C subfamilies. a Correlation of log2 fold enrichment relative to the random genomic background between two ESCs from different sources. ERV/LTR subfamilies are included. Log2 fold enrichment was computed as the number of instances overlapped with the actual ATAC-seq or DNase peaks versus 1000 randomized peaks (see Methods). *R^2^* and *p* values were computed by the linear regression model. b Log2 fold enrichment of the chromatin accessibility and H3K27ac mark per TE subfamily in ESCs relative to NPCs. c Log2 fold enrichment of candidate subfamilies (Supplementary Fig. 5B) with increased or decreased accessibility and H3K27ac activity during the differentiation from ESCs to NPCs (72 hours). d Log2 fold enrichment of candidate subfamilies (Supplementary Fig. 5B) across five cell types. Chromatin accessibility and H3K27ac were analyzed per cell type. e Log2 fold enrichment of each KRAB-ZNF overlapped with each MER11 subfamily in HEK293T cells. Available KRAB-ZNF binding sites in HEK293T cells (Imbeault, Helleboid, and Trono 2017) overlapped with a minimum of 20 instances and more than two-fold enrichment related to the random genomic background with *p* value ≤ 0.05 are highlighted. *P* values were computed using the same permutation test. We consistently found the enrichment of ZNF525, ZNF808, ZNF440, ZNF433, and ZNF468 in these MER11 subfamilies. Moreover, we newly identified the enrichment of ZNF33A and ZNF611 in MER11A subfamily. f Proportion of instances per MER11 subfamily overlapped with KAP1 in HEK293T cells. 73.2% of MER11A, 68.2% of MER11B, and 42.4% of MER11C are also bound by KAP1, confirming their potential repression in HEK293T cells. g Highest and lowest proportion of peaks-associated instances among MER11 new subfamilies or original subfamilies per epigenetic mark.

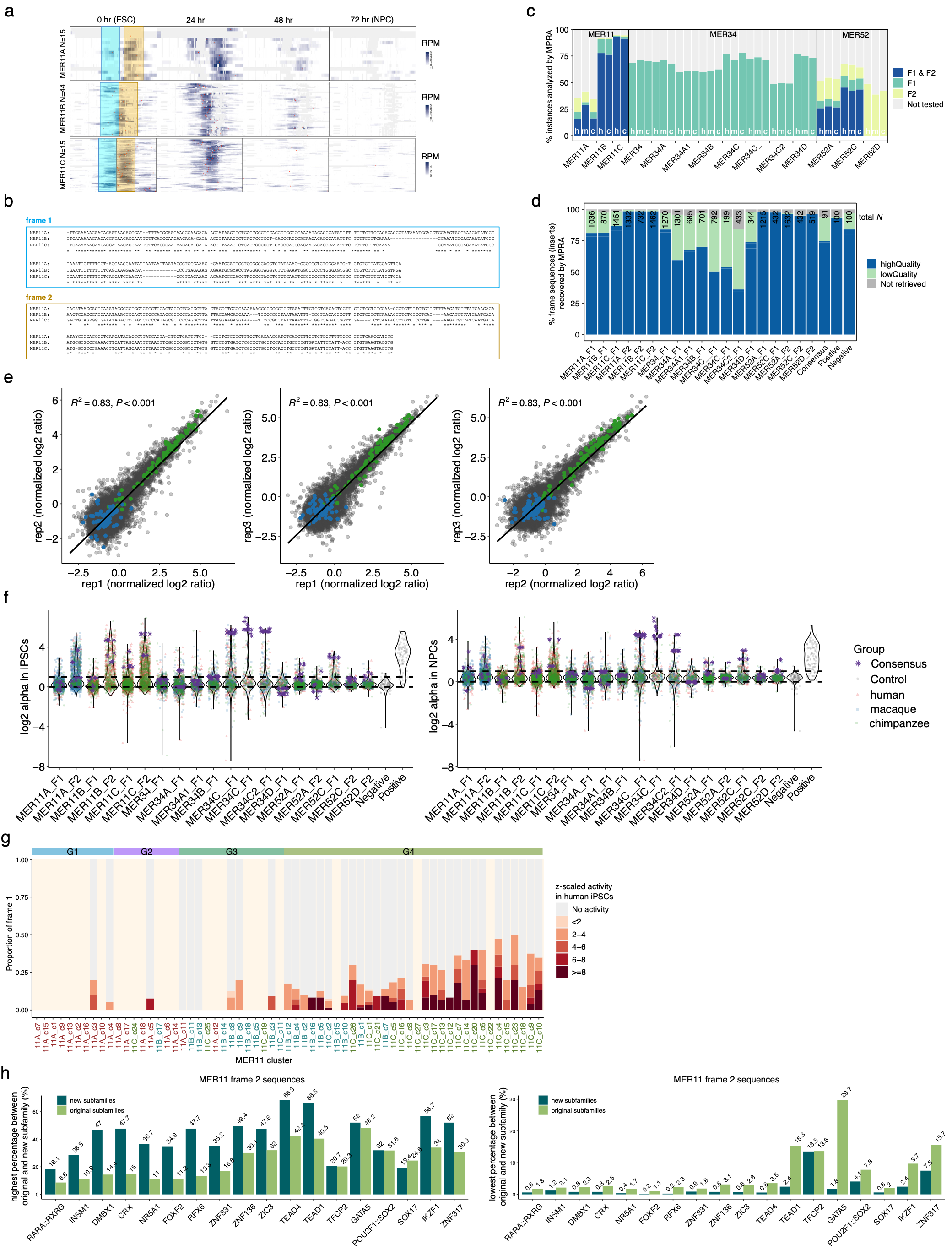

#### Supplementary Figure 6 MPRA activity of MER11, MER34, and MER52 frame sequences in human iPSCs and differentiated NPCs. a ATAC-seq read distribution (read per million) on each accessible MER11A/B/C instance along each consensus sequence. Frame 1 and frame 2 regions are highlighted in light blue and brown. b Multiple sequence alignment of extracted MER11A/B/C frame 1/2 consensus sequences. ClustalW2 with default parameters (<https://www.ebi.ac.uk/Tools/msa/clustalw2/>) was used for the analysis. c Proportion of genomic instances (≥ 200 bp) from each of MER11, MER34, and MER52 subfamilies analyzed by MPRA. Human (hg19), chimpanzee (panTro4) and macaque (macFas5) genomes were analyzed. d Proportion of MER11, MER34, and MER52 1/2 frame sequences successfully retrieved by MPRA experiment. Number of consensus sequences refers to the sequences different from other frame sequences. High-quality inserts refer to examined frame sequences associated with ≥ 10 barcodes in two or more DNA libraries. Low-quality inserts refer to frame sequences associated with < 10 barcodes in less than two DNA libraries. Not retrieved inserts refer to frame sequences that fail to be associated with any barcodes in three replicates. e Correlation of normalized log2 RNA/DNA ratio in human iPSCs between three replicates. Normalized RNA/DNA ratio was achieved by MPRAflow (see Methods). Positive and negative controls are in green and blue color. f Violin plots of log2 alpha values per subfamily in human iPSCs and NPCs. Alpha value was computed by MPRAnalyze (see Methods). Dotted lines indicate the 0 and 1 log2 alpha values. Consensus sequences with different nucleotides for each ambiguous nucleotide position are included. g Proportion of active MER11 frame 1 sequences per cluster. Clusters with less than 10 instances measured by MPRA are excluded and shown as the background color (light pink). h Highest and lowest proportion of instances containing each motif amongst new subfamilies or original subfamilies.

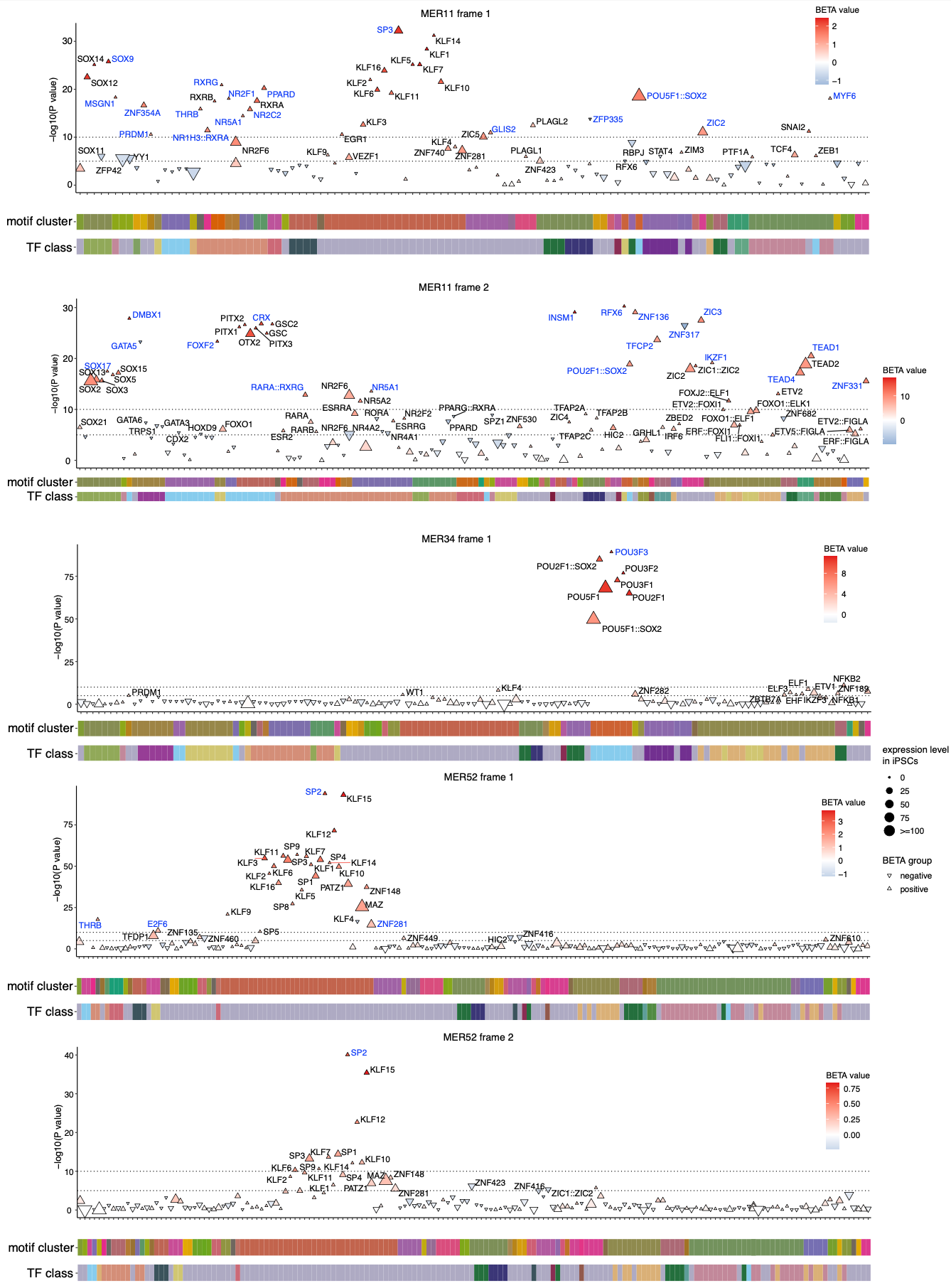

#### Supplementary Figure 7 Motifs associated with the MPRA activity. Motifs significantly associated with the MPRA activity are highlighted (-log10 *p* value ≥ 5) and the top selected motifs amongst each motif group are highlighted in blue. *P* value refers to the significance and was computed using plink2 (see Methods). BETA value refers to the effect size. Motif cluster and TF class information could be found at <https://jaspar.genereg.net/matrix-clusters/vertebrates/?detail=false>.

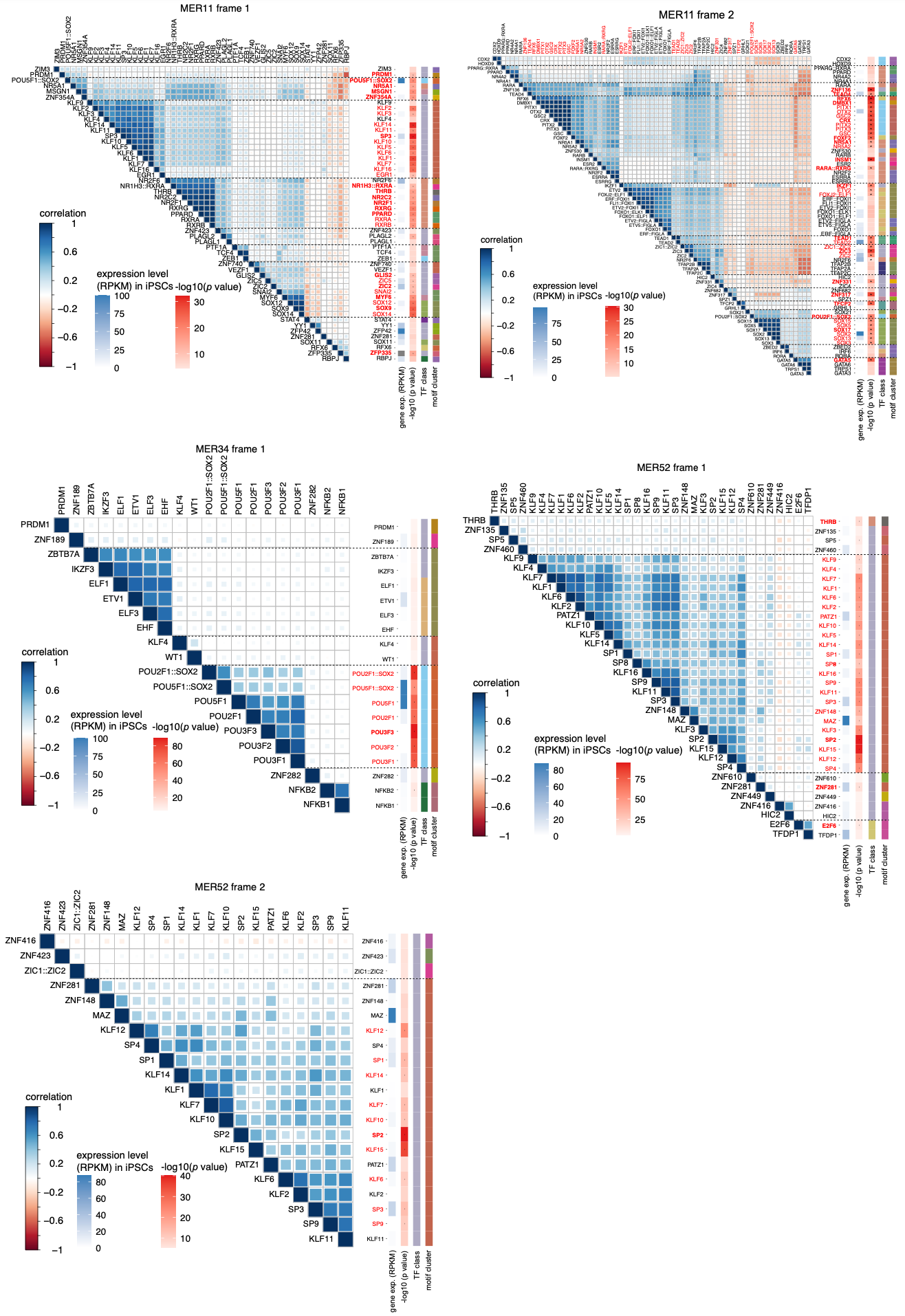

**Supplementary Figure 8** Correlation amongst candidate motifs per frame region. Motifs with *p* value ≤ 1 × 10^-5^ are selected as the candidates. Motifs with *p* value ≤ 1 × 10^-10^ are highlighted in red. Dotted lines separate the different groups of strongly associated motifs. Top candidates within each group are highlighted in bold.

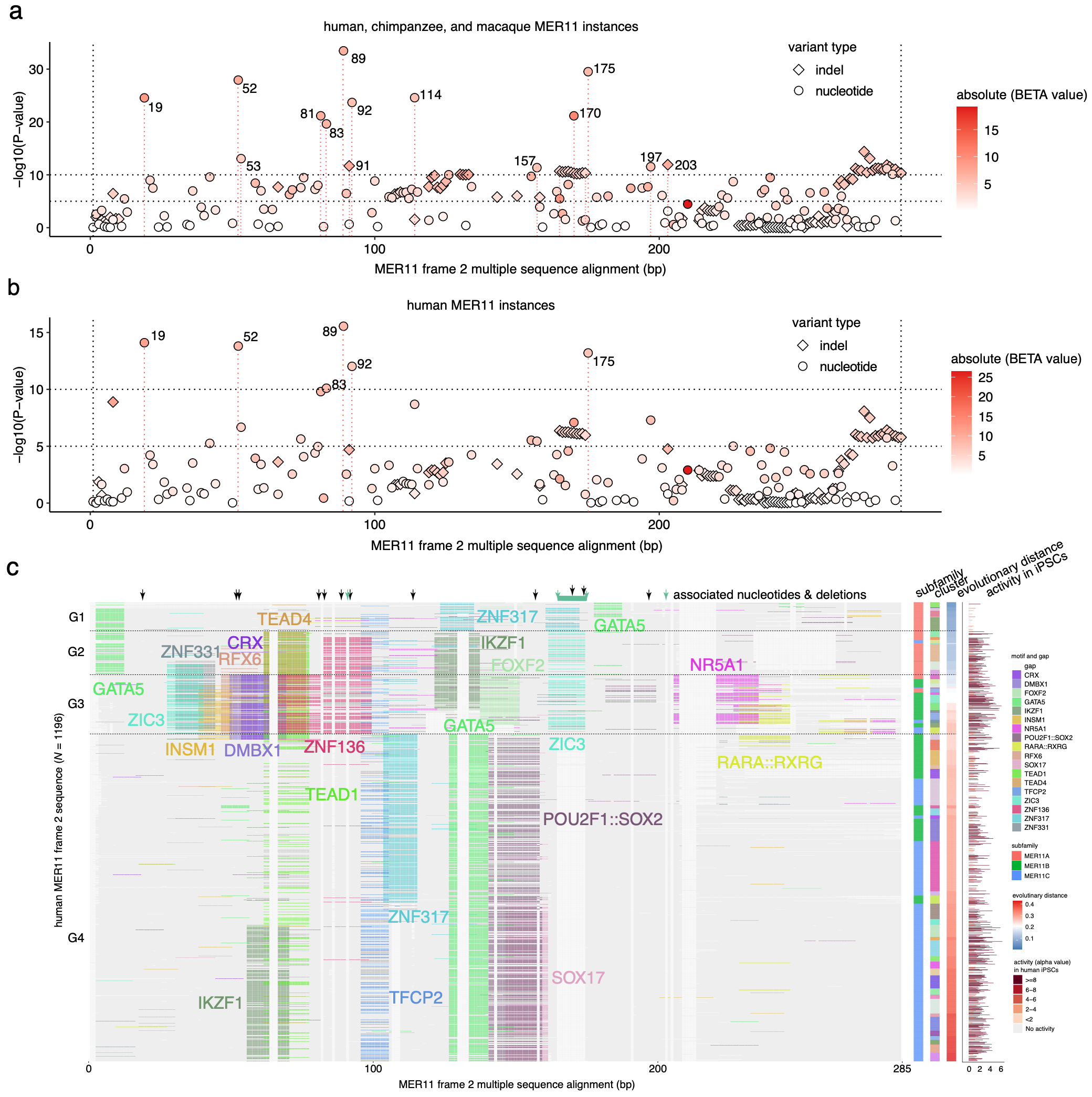

#### Supplementary Figure 9 Nucleotides and motifs associated with the MPRA activity. a Nucleotides significantly associated with the MPRA activity amongst all MER11 frame 2 genomic sequences. Nucleotide positions refer to the locations along the multiple sequence alignment. Alleles of each indel and nucleotide mutation are shown separately. b Nucleotides significantly associated with the MPRA activity amongst human MER11 frame 2 sequences. Nucleotide positions refer to the locations along the multiple sequence alignment. Alleles of each indel and nucleotide change are shown separately. c Positions of associated nucleotides and motifs on the human MER11 frame 2 multiple sequence alignment. Associated nucleotides detected using the human, chimpanzee, and macaque sequences are shown. Detected motifs and nucleotides on the multiple sequence alignment of human sequences is shown as an example. Black and green arrows indicate the positions of strongly associated nucleotide mutations and indels.

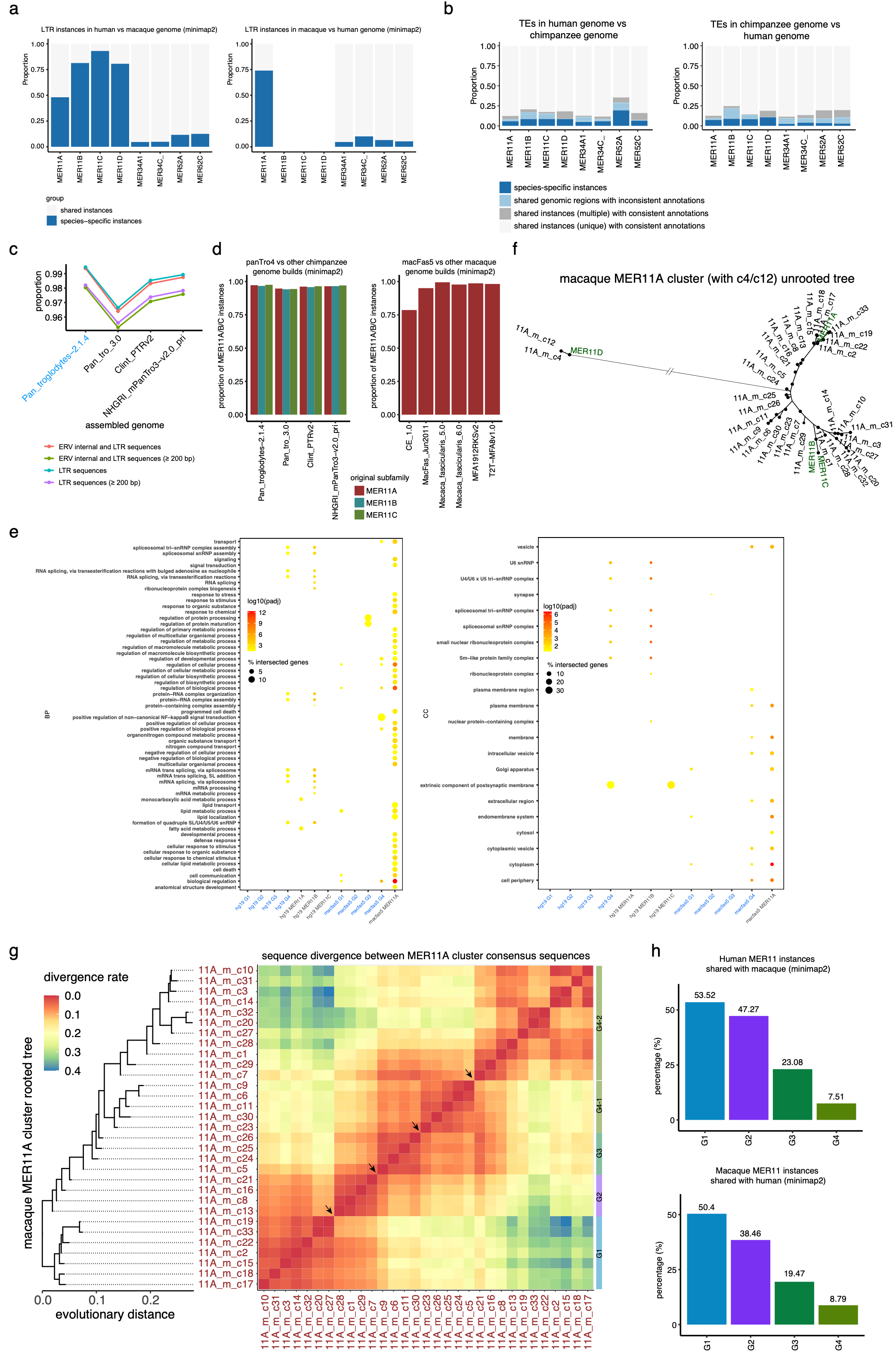

#### Supplementary Figure 10 Evolution and MPRA activity of macaque MER11A subfamily. a Conservation and annotation of MER11 instances in human versus chimpanzee based on minimap2 alignment (see Methods). b Conservation and annotation of MER11 instances in human versus chimpanzee based on liftOver and RepeatMasker. Instances intersected with regions annotated as the same or different subfamilies in another species are shown separately. Instances intersected with unique or multiple regions are also shown separately. c The conservation of annotated panTro4 LTR sequences in other Pan Troglodytes genome builds. LTR/ERV subfamilies (with or without internal sequences) were calculated separately. d Conservation of MER11 instances in human versus chimpanzee based on minimap2 alignment. e Biological process (BP) and cellular component (CC) analysis of MER11 original and new subfamilies in human and macaque. The closest gene to each MER11 instance was achieved using Bedtool2 *closest* function. The distance between MER11 and cloest genes within 100 kb was kept. G:profiler online tool was used for the gene enrichment analysis per original/new subfamily. f Unrooted tree of all MER11A cluster consensus sequences. MER11A/B/C/D subfamily consensus sequences are highlighted in green. g Rooted tree and divergent rates of macaque MER11A cluster consensus sequences. 11A_m_c4/c12 were excluded. New subfamilies are determined as we previously described (see Methods). Due to the large number of macaque MER11A instances, new subfamilies with a minimum of 50 were kept. Arrows indicate the boundaries between them. Macaque MER11_G4-1/G4-2 are defined according to their sequence similarity with human MER11_G4 cluster consensus sequences.

**h** Proportion of MER11 instances per new subfamily shared between human and macaque. The shared instances were identified by minimap2 alignment.

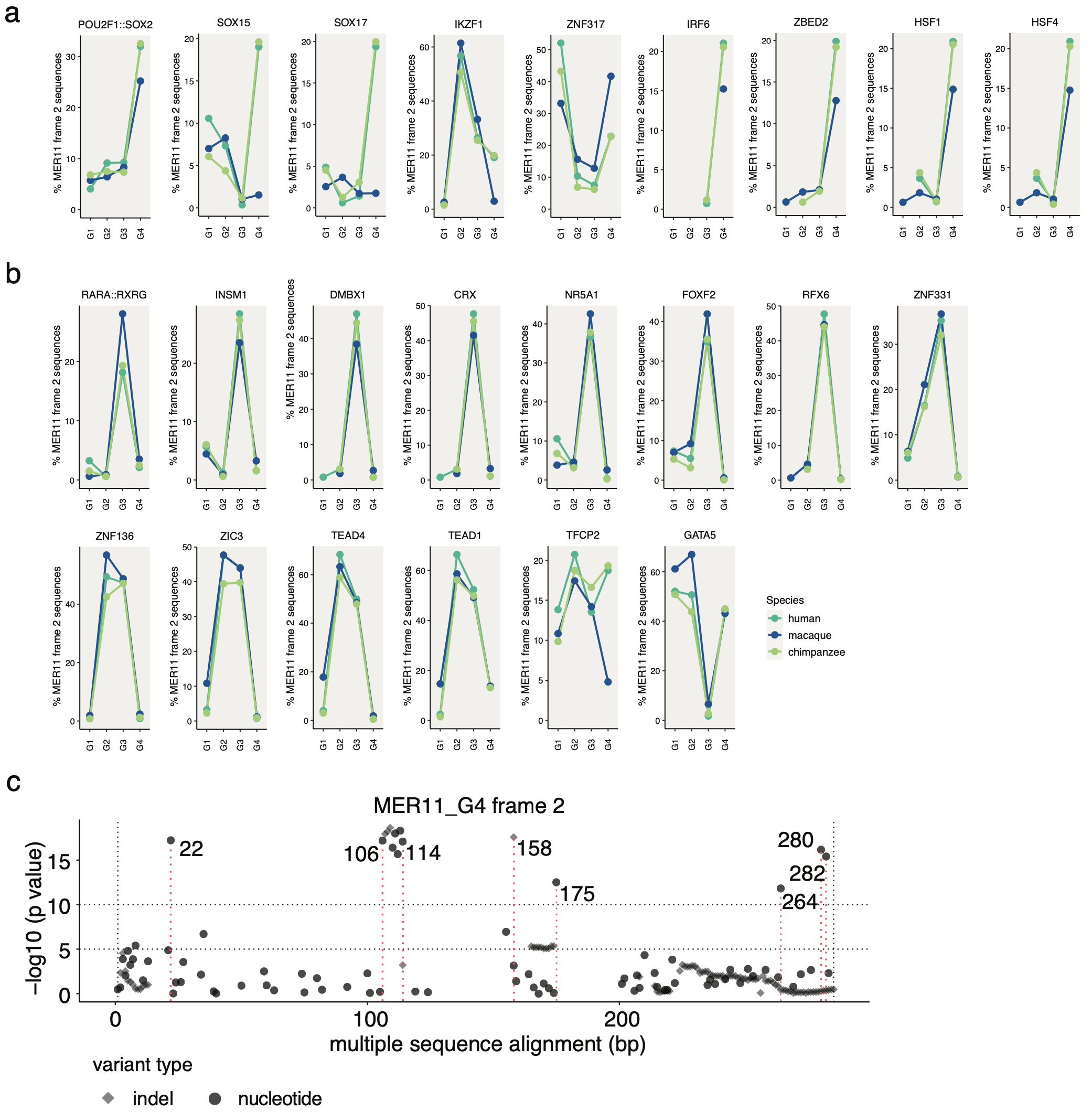

#### Supplementary Figure 11 Nucleotides and motifs associated with the MPRA activity between species. a Proportion of frame 2 sequences containing each candidate species-specific motif (e.g., SOX15) across human, chimpanzee, and macaque new subfamilies. b Proportion of frame 2 sequences containing each associated and none species-specific motif across human, chimpanzee, and macaque new subfamilies. c Nucleotides associated with the MPRA activity across all MER11_G4 frame 2 sequences. Human, chimpanzee, and macaque sequences were included to increase the detection power.

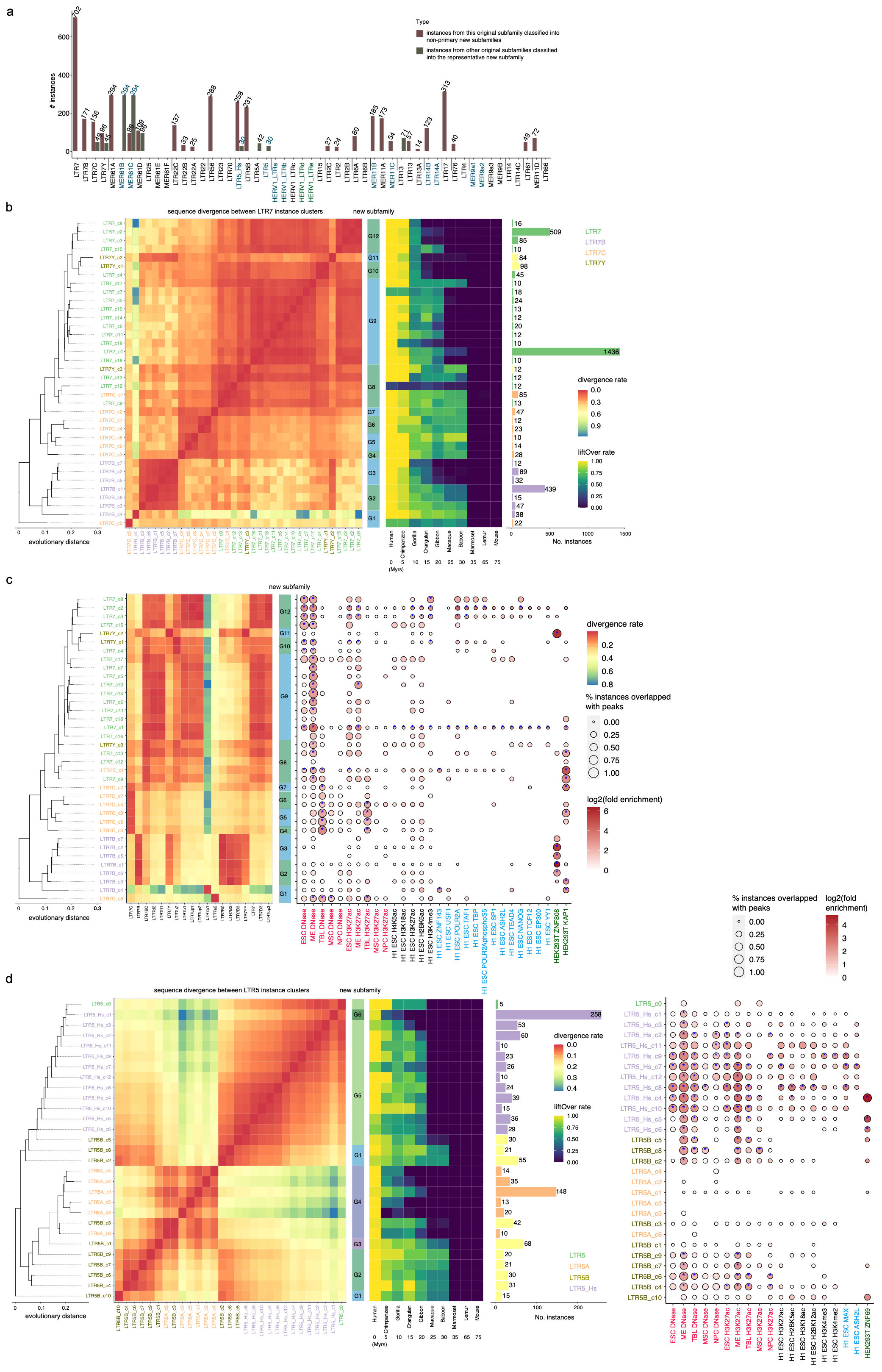

#### Supplementary Figure 12 Phylogenetic and epigenetic analysis of LTR5 and LTR7 subfamily groups. a Number of misannotated instances corrected by defined new subfamilies. Subfamilies are ordered by the number of new subfamilies per subfamily group. The new subfamily containing the most instances (top) is used to represent each subfamily. Subfamilies with the same representative new subfamilies are highlighted in blue or green. Number of misannotated instances counted in two subfamilies is highlighted in blue. b LTR7 subfamily group rooted tree and the sequence similarity between clusters. c Epigenetic profile of every LTR7 cluster and new subfamily and the sequence similarity with publicly available subfamily consensus sequences. Subfamily consensus sequences were downloaded from the DFAM database. Epigenetic marks overlapped with a minimum of five instances for small clusters (< 100 instances) and 20 instances for large clusters (≥ 100 instances) were kept. Permutation was used to compute the *p* values. Significantly enriched (log2((actual counts+1)/(mean shuffled counts + 1)) ≥ 1 and *p* value ≤ 0.05) clusters relative to 100 random genomic controls are highlighted. d LTR5 subfamily group rooted tree and the sequence similarity and epigenetic profile of every cluster.

### Supplementary Tables

**Supplementary Data 1** Genome assemblies used for the orthologous analysis across primate species.

**Supplementary Data 2** Summary of available macaque fascicularis (crab-eating macaque) genomes.

**Supplementary Data 3** List of refined annotations of 53 simian-enriched LTR subfamilies. Instances shorter than 200 bp were also annotated by *Blastn-short* against every cluster consensus sequence from each subfamily group. The top target cluster consensus sequence with a minimum of 50% alignment length versus the instance length was kept. Instances that failed to be annotated were classified as the unannotated group. We also re-annotated 2,926 instances from analyzed 53 subfamilies that were shorter than 200 bp through Blastn against each cluster consensus sequence and 2,055 of them were classified into different clusters and new subfamilies.

**Supplementary Data 4** Summary of MER11, MER34, and MER52 sequences submitted to MPRA.

**Supplementary Data 5** Characteristics of examined MER11, MER34, and MER52 frame sequences by MPRA. Subfamily consensus sequences, and positive and negative control sequences were also included. Computed alpha values, *z*-scaled alpha values and corresponding *p* values in iPSCs and NPCs were also included.

**Supplementary Data 6** Summary of available Pan Troglodytes (chimpanzee) genomes.

**Supplementary Data 7** List of genes closest to the human and macaque MER11_G1/G2/G3/G4 instances.

**Supplementary Data 8** List of new subfamilies and clusters amongst 53 simian-enriched LTR subfamilies. The number of instances, branch length (bootstrap value) to the selected root, and the liftOver rate to the macaque genome per cluster were included.

**Supplementary Data 9** Epigenetic profile of every new subfamily amongst 53 simian-enriched LTR subfamilies.

**Supplementary Data 10** List of primers used for lentiMPRA.
